# Detecting the body’s reproductive hormonal brake against tissue overgrowth: micrin/SgII-70

**DOI:** 10.1101/2024.10.01.616062

**Authors:** John E Hart, Keith G Davies, Christopher R Mundy, Aurelia C Hart, David R Howlett, Russell P Newton

## Abstract

A new humoral factor has been detected, within a project aiming to disclose the body’s reproductive hormonal brake against tissue overgrowth, micrin (‘my-crin’). It is hypothesized that micrin braking, gonadal and hypothalamic, is lifted at puberty and wanes with age, bringing on prostatic enlargement and cancers. Factor purification has involved fractionation of ovine and bovine ovarian follicular fluid and blood plasma and serum, with evaluation via rat bioassays in vivo and in vitro. Analysis averse, the molecular effector provided a chemical conundrum. Evidence from mass spectrometry (MS) has been problematic, with spectra from MALDI-TOF, the only productive MS modality, confused by a target polypeptide exhibiting artefacts during processing, storage and MALDI set-up in terms of C-terminal truncation and water losses, together with N-terminal grand fragmentations and dimerizations (factor fragments doubled up). Evidence from chemical sequencing of amino acid (aa) residues was likewise baffling, but consistent with a spiralised depolymerisation within the Edman reaction chamber of a unitary polypeptide. Data decryption has overcome molecular intractability, supported by the results of tryptic digestion and epitope mapping using immunohistochemistry (IHC). The detected inhibitory factor is sleuthed to relate to secretogranin II (SgII), the neurosecretory prohormone, in the form of a secreted acidic 70-aa polypeptide derivative called here SgII-70 (‘sig two seventy’). The product of peptide splicing, micrin/SgII-70 is potentially an amphipathic molecular ampersand (&), with ends entwined (via salt bridging), the knotty totality compromising Edman and MALDI analyses and molecular modelling whilst conferring protease and heat resistance. The two ends of the factor are conceptualised as binding at the same time to a dimeric cellular receptor, to stoichiometric effect, providing fewer smaller cells within a counting mechanism of tissue-mass regulation. Hexapeptide mimetics simulating both ends of the hormone together have been demonstrated in different species and settings, in the cause of antiorganotrophism (tissue reduction) and reproductive modulation. Therapeutic exploitation beckons for mimetics of the body’s hormonal brake, micrin/SgII-70, in tissue overgrowth conditions such as endometriosis, PCOS, BPH and cancer, and in infertility.

## Background

There is a paradoxical increase in pituitary weight in both female rats and mice in the first few weeks after ovariectomy, seemingly indicating the abolition of a net inhibitory ovarian influence, the established hypophysiotrophic effect of oestrogens notwithstanding (Hart, 1990a). In the male rat, castration also leads to pituitary hypertrophy, an effect that can be reversed by an aqueous testicular extract but not by a lipid testicular extract (steroid rich) or by testosterone alone (McCullagh, 1932). Exogenous oestrogens are the most organotrophic substances known, enlarging organs visceral (e.g. heart, liver, kidneys), endocrine (e.g. pituitary, adrenal, ovaries) and reproductive (e.g. uterus) (Hart, 1990a). The organotrophic effects of exogenous oestrogens are not seen in hypophysectomized female rats, suggesting an indirect effect via the hypothalamic-pituitary axis (Hart, 1990b). The organotrophic oestrogen effect in female rats is more marked in ovariectomised animals than in intact ones (Marlow, 1939; Matthews et al, 1942; Smith, 1947; Hart, 2014). This observation itself speaks of gonadal inhibition. It has been hypothesized that the gonadal inhibitory influence is stimulated by antioestrogens, accounting in part for the antiorganotrophic activity of these compounds in rodents and their therapeutic usefulness in humans (Hart, 2014). The antioestrogen clomiphene, for example, reduces the size of the rat pituitary, liver, spleen, kidneys, heart, uterus and ovaries in adults whose body weights are unchanged (Hart, 1990b). The antioestrogenic blockade of the organotrophic effects of exogenous oestrogens is blunted by prior removal of the ovaries.

Brandishing Occam’s Razor, the foregoing and other pieces of evidence led to the suggestion that the body has a unitary hormonal reproductive brake against tissue overgrowth, non-steroidal in character, which brake is lifted to initiate puberty and the assumption of adult stature and whose action wanes with age, leading to a rise in cancers and the enlargement of the prostate (Hart, 2014). This hypothetical hormonal factor, dubbed ‘micrin’ (pronounced *my-crin*; Greek, *small*, as in small organs), is conceived as being a potentially novel inhibitory effector in the hypothalamic-pituitary-gonadal component of the ‘organotrophic system’ of internal size regulation. The hunt for a hormone (long-distance chemical messenger) described here implicitly tests alternative hypotheses such as ‘factor non-existence’ and ‘existing factor with newly recognized antiorganotrophic activity’. The narrative form of the paper arises from the integration of a body of novel data (notably MS) with disclosures from prior publications and dispersed in the patent literature.

### Purification

The search for the molecular explanation of an inhibitory gonadal hormonal influence on mammalian tissue masses featured bioassay guided fractionations involved ovarian follicular fluid, ovarian venous plasma and systemic blood plasma and serum mainly from ewes (*Ovis aries*), but also cattle (*Bos taurus*) and pigs (*Sus scrofa domesticus*), with testing in bioassays in vivo and in vitro in the rat (*Rattus norvegicus*).

Fractionation by the canonical method involved spin filtration (sizing membranes) for separation and concentration plus gel filtration (size-exclusion chromatography), with anion exchange chromatography (‘anionex’, using a positively charged resin) for final purification (Beale, 1969). Mass analysis involved MALDI-TOF MS (Matrix-Assisted Laser Desorption/Ionization - Time-of-Flight Mass Spectrometry, hereafter ‘MALDI’; positive ion, linear mode unless otherwise specified, delayed extraction typically of 350 nsec, low mass gate usually set at 1000 Da) and SDS-PAGE (Sodium Dodecyl Sulphate - Polyacrylamide Gel

Electrophoresis, hereafter ‘gel’; mainly 1D). The purification method was developed at The Babraham Institute, Cambridge, UK, hereafter ‘Babraham’ (see Babraham Method in Supplementary Information 1, Candidates & Purifications = S1 of 7). Anti-organotrophic activity was assessed in vivo by shrinkage (weight loss) of internal organs over four days in the young adult female intact rat and in vitro by inhibitory effects on the viability of rat bone marrow cells (BMCs) in primary culture (S2 Assay Methods & Results). In the BMC assay, bone marrow in semisolid form is centrifuged from sections of femoral and tibial bone and cultured in media (Dobson et al, 1999), to provide an organ-like mixed-cell population including haematopoietic stem and progenitor cells and mesenchymal stromal cells such as fibroblasts and pericytes.

An internal-organ-reducing activity has been tracked through successive stages in the purification of sheep’s blood (Hart, 1999), using the rat organometric assay in vivo for guidance, to maximum purity represented by anionex fractions. An organotrophic pattern of ‘pituitary down, adrenals up’ is claimed to be a micrin signature (Hart, 2014). This is seen in Fig. 1 in rats exposed to anionex HPLC fractions of ovarian venous plasma from ovary-intact (OV-INTACT) sheep when compared to rats receiving such fractions from the jugular vein plasma of ovariectomised (OVX) ewes. This result has been replicated (S2). The surprising pattern of ‘pituitary down, adrenals up’, though not invariably evoked, has been recorded in other circumstances: (a) when using steroid-depleted aqueous beef testicular extract in male intact rats, in which ‘prostate down’ is also seen (Vidgoff and Vehrs, 1940); (b) in male intact rats receiving a 0.1-0.2 M NaCl anionex FPLC fraction of ovarian venous plasma previously subjected to spin and gel filtration to provide a 10-20 kDa fraction (S1 Babraham Method), in which the relative organ weights vs controls after a few days were pituitary (−2.3%), adrenals (+9.7%) and prostate (−23.4%) (Hart, 1999); (c) in female rats treated with clomiphene (Hart, 1990b), a putative micrin stimulator (Hart, 2014); (d) in the same experiment, where clomiphene blunted oestrogen-induced pituitary and other organ enlargements but exaggerated an oestrogen-induced adrenal enlargement; and (e) in rats exposed in the standard organometric assay to four daily doses of anionex fractions (S1 Babraham Method) of ovarian venous plasma from intact sheep which had been treated with either clomiphene (test) or saline (control), with the result that the relative organ weights of the test versus control rats varied as follows: pituitary −8.4%; heart −3.9; liver −9.5%; adrenals +4.1%; kidneys −6.0%; uterus −27.4%; ovaries −6.4%; and spleen −1.1%. (Further examples are depicted in S2.) The same directionality difference is seen in young rats after ovariectomy, with pituitary and other organ weights above those of intact animals, adrenals below (Freudenberger and Howard, 1937). Antihypophysiotrophic and broadly antiorganotrophic, micrin is yet adrenotrophic.

**Figure 1.**
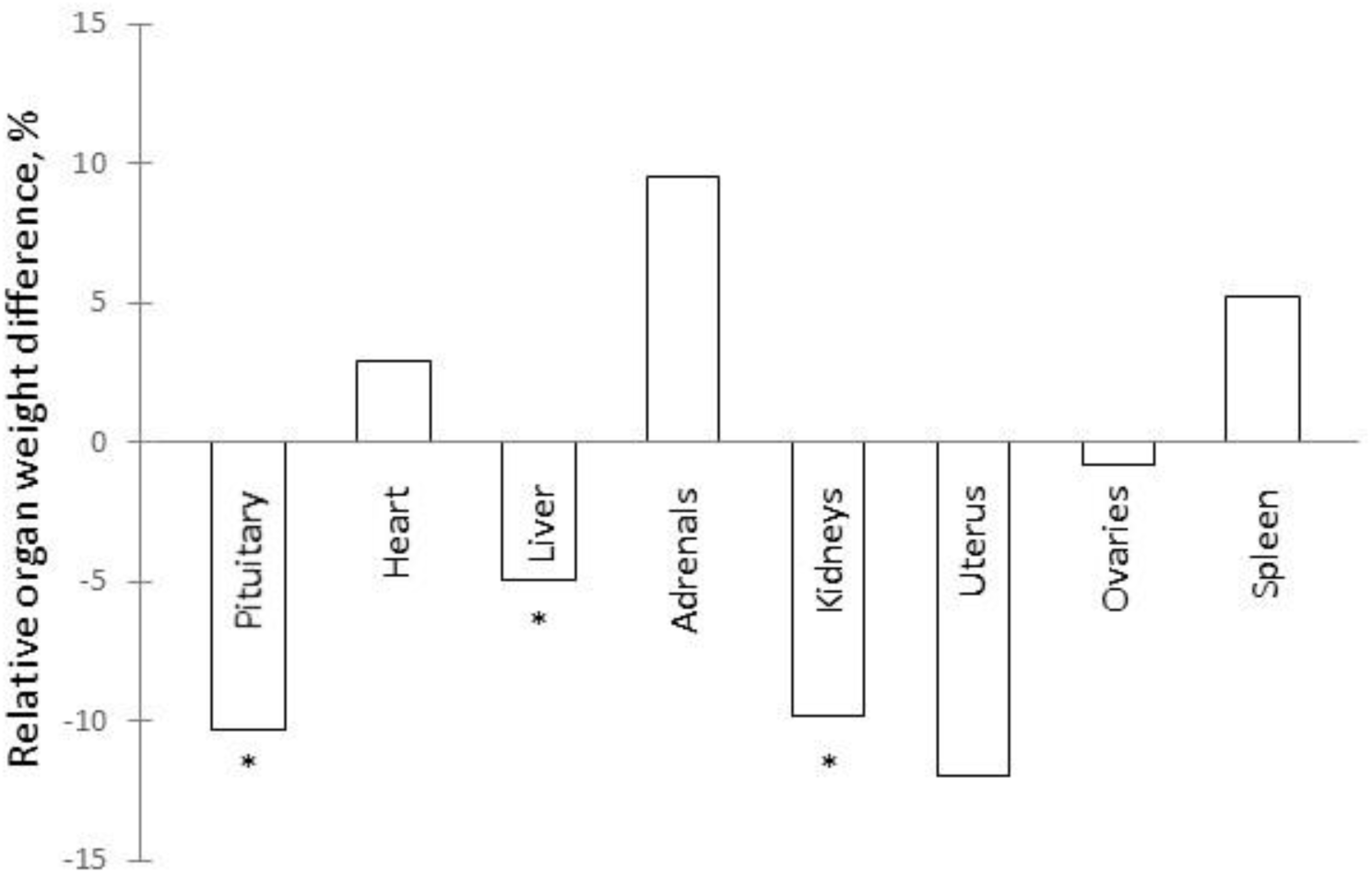
Organometric effects in rats in vivo of plasma anionex fractions from OV-INTACT vs OVX sheep. Post-mortem organ weights expressed as a percentage of body weight for two groups of seven intact adult female Sprague Dawley rats. The zero baseline is provided by rats receiving for four days by intraperitoneal injection a fraction (spin and gel filtered to 10-20 kDa, eluting in HPLC anionex at ∼0.2 M NaCl; S1 Babraham Method) of the jugular vein plasmas pooled from two OVX sheep. The other group of rats received the same fraction in the same amount from ovarian venous plasmas pooled from six OV-INTACT sheep. No significant body weight changes were seen to confound the picture of organ shrinkage with adrenal enlargement. * = P<0.05, one-tailed t test. (N.B. OV-INTACT versus PBS controls, adrenals +8.53%, significant: S2. The adrenal enlargement in these rats is non-artefactual.)

In the rat study in vivo that furnishes Fig. 1, gross abnormalities were absent and a histological investigation confirmed an apparent lack of toxicity, in line with a lack of behavioural deficits. The organ mass alterations reported here and previously (Hart, 1999) would appear to be due to cell shrinkage, reduced cell numbers by apoptosis and suppression of angiogenesis, according to the results of assays in vitro (S2). In summary, fewer smaller cells.

In the standard BMC cell death assay in vitro use was made of jugular vein whole plasma from OVX sheep (n = 2) and ovarian vein whole plasma from OV-INTACT sheep (n = 22), the neat plasmas being putatively micrin-minus and micrin-plus, respectively. Cell survival after 48h (assessed by Alamar Blue) was normalised against one of the OVX samples as 100%, with the other at 98%. For the 22 OV-INTACT samples the mean +/– SEM was 25.7 +/– 4.7% (P<0.0001, two-tailed t test, significant). Cell shrinkage was evaluated in the same system using Magiscan image analysis, though the results were obvious to the microscope-aided eye Hart, 2000). The cell cross-sectional areas (all cells in 10 microscopic fields at x40 magnification, mean +/– SD, arbitrary units) after 48h were: saline control, 0.67 +/– 0.38; plasma from OVX sheep, 0.78 +/– 0.55 (the mean differing from saline by +16%, ns by t-test); and plasma from OV-INTACT, 0.47 +/– 0.23 (–30%, P<0.005). The differences will have been greater of course in volume terms. In another study series selected anionex fractions of sheep serum produced a significant increase in apoptotic cells, as judged by Annexin V and TUNEL assessments, without an increase in necrotic cells, suggesting a physiological effect rather than toxicity (S2).

From column and solvent studies used in conjunction with assays of BMC viability it was judged that the growth inhibitory agent (singular for parsimony) did not have the characteristics of a lipid or a carbohydrate (Wiles, 2002). (Delipidated sheep serum samples still contained the main candidate molecule from MALDI, to be described shortly.) The molecular weight was probably ∼10 kDa and the factor was possibly polar. Charcoal reduced bioactivity in vitro, probably indicating a charged molecule. A negative charge was anyway implied by the success of the purification campaign in generating anionex fractions that shrank multiple organs in vivo (Fig. 1). Peptidic character was suspected. In line with this, the activity in vitro was reduced by the proteolytic enzyme trypsin and, separately, heat, as will be described later. In terms of retention of bioactivity in vitro, plasma from blood anticoagulated with EDTA proved to be a better purification feedstock than serum or heparinised plasma (S1). As EDTA inhibits proteolytic enzymes, this is consistent with micrin having a proteinaceous character. Loss of bioactivity in vivo and in vitro has been seen after storage short-term (a few days), possibly suggestive of susceptibility to protease activity or more likely pure-form lability (i.e. instability). Lyophilization did not conquer apparent lability, instead exacerbating loss of activity after storage (S2 Fig. 23).

### Candidates

During an extensive purification campaign, a dozen candidates were seen for the sought-for tissue-mass inhibitory reproductive hormonal influence in mammals (S1). To summarise the bioassay-guided physicochemical fractionation campaign, two things emerged: (i) ‘**<u>Candidate</u> <u>7500’</u>**, a group of peaks (ions) in the mass-to-charge (*m/z*) range 7-8000 (Figs. 2-6) seen in MALDI (at Babraham, unless otherwise stated); and (ii) ‘**<u>EPL001’</u>**, an Edman machine N-terminal sequence of 14 aa residues, MKPLTGKVKEFNNI (S1 Table 1; likewise obtained at Babraham), relating to an ovine blood plasma fraction containing Candidate 7500.

**Figure 2.**
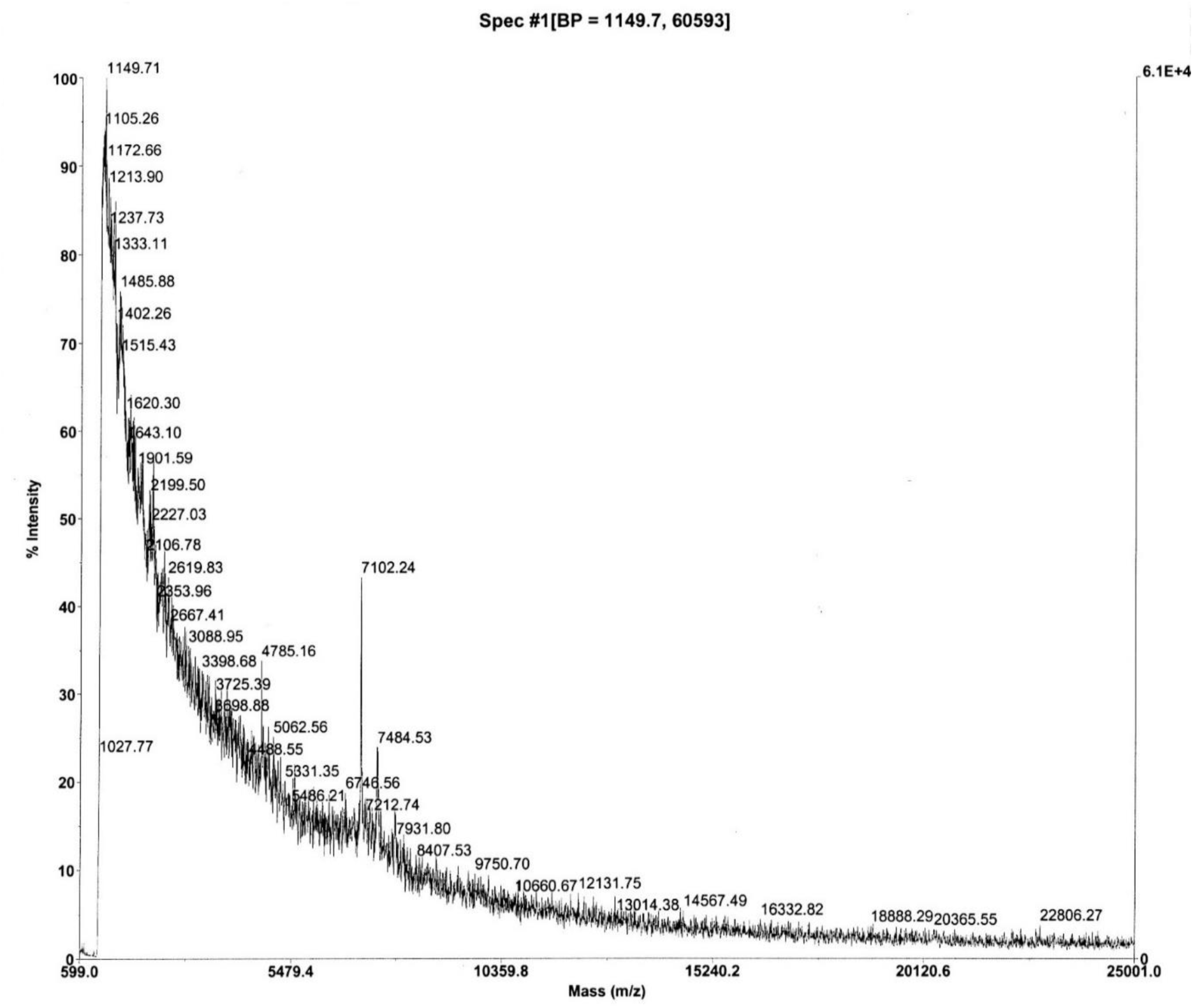
MALDI mass spectrum of ovine ovarian follicular fluid. Anion exchange Fraction 27 (i.e. late eluting), after spin and gel filtration. Matrix sinapinic acid, calibration against carbonic anhydrase (Babraham MS, S3). For Candidate 7500 MS peak analysis in regard to Figs. 2-5 see Table 1. The peak at *m/z* 4785 is analysed to be a **42mer** N-terminal fragment of sSgII-70 and the 1149 base peak (‘BP’ in header) an **11mer**. (See S3 Figs. 1 & 2 for equivalent bovine and pig spectra.).

**Figure 3.**
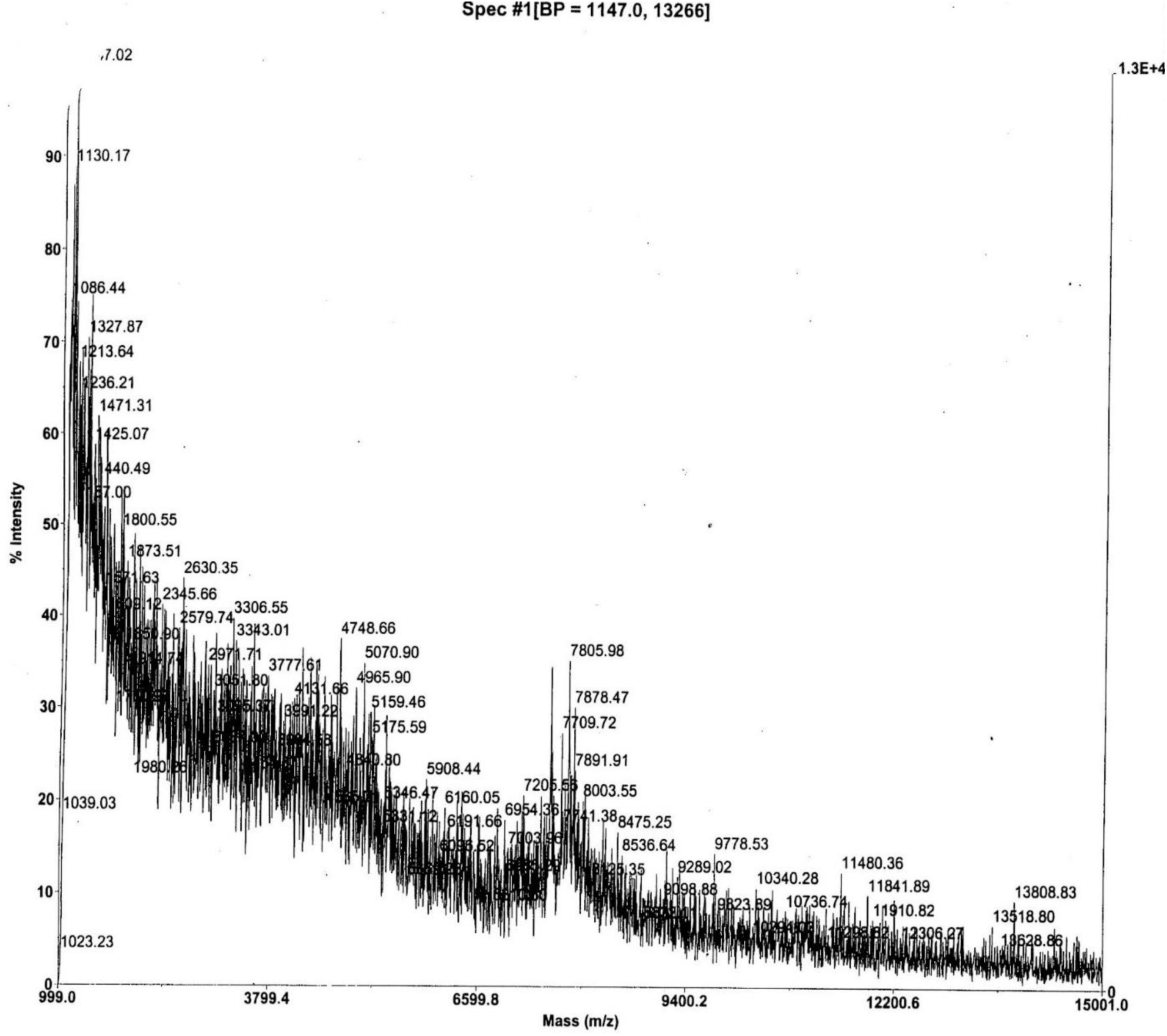
MALDI mass spectrum of ovine jugular vein EDTA plasma anionex Fraction 29. The Beale 4 residues were determined in this material by Edman sequencing from a gel lacking in visible bands but excised anyway at ∼7 kDa. The Beale 4 aa correspond to EPL001 residues 3, 8, 11 and 12, without EPL001’s 10 other aa being discernable, thus: _1_MK**<u>P</u>**LTGK**<u>V</u>**KE**<u>FN</u>**NI_14_ (S1 Table 1, SEQ ID NOs: 3 & 4). MALDI matrix and calibration as Fig. 2. The ion at *m/z* 7805 is an integer match to a predicted sSgII-70 **68mer**. A peak without mass annotation is ‘c7500’. Minor peaks at 5070 and 4748 are analysed to be **45mer** and **43mer** fragments of sSgII-70, respectively. The base peak is missing from this photocopy record but is 1147 (‘B.P.’ in header), an sSgII-70 **11mer**.

**Figure 4.**
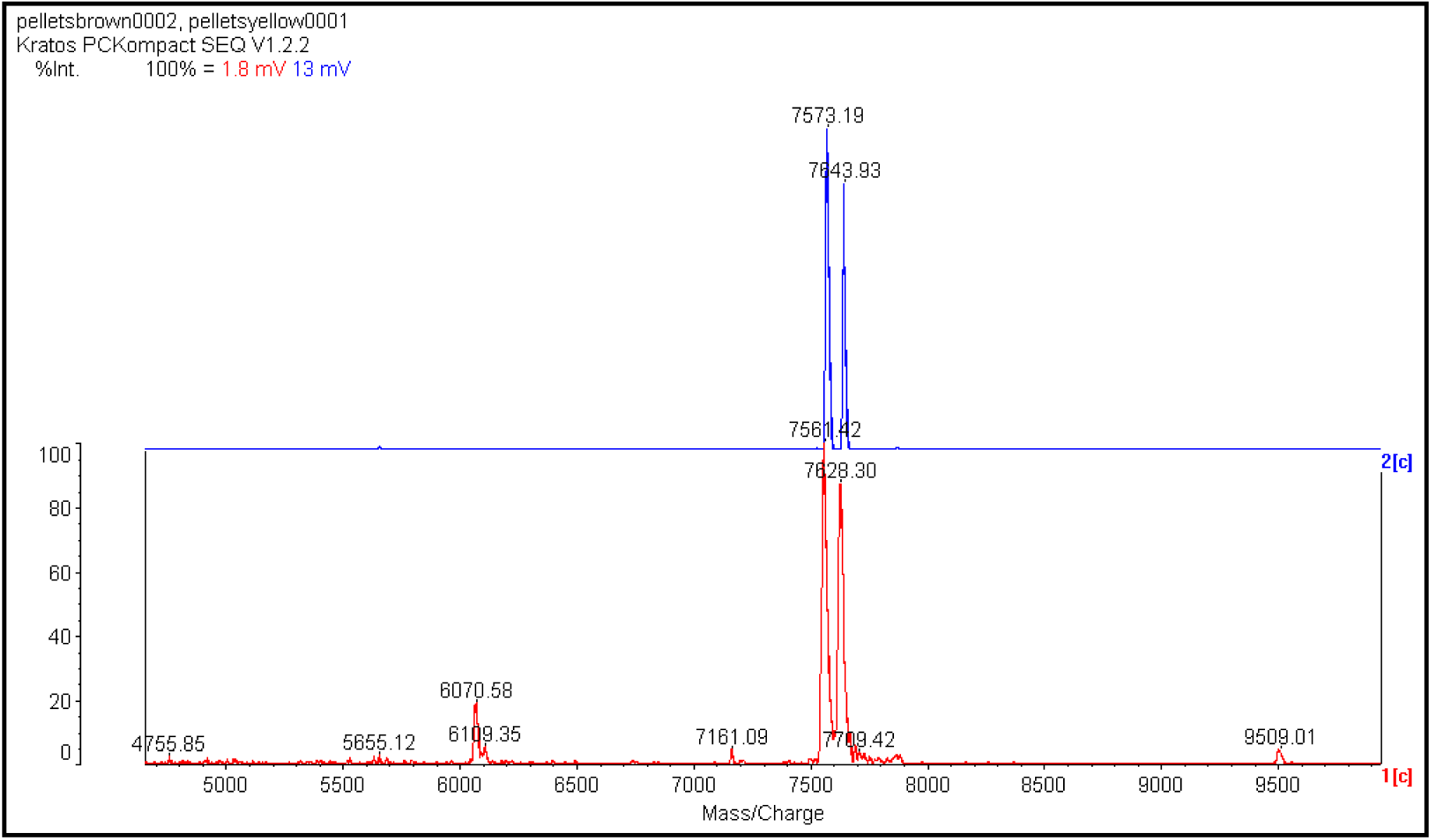
MALDI mass spectra of a solid ‘upstream precipitate’ that formed on the 3 kDa membrane during a 3-30 kDa ultrafiltration of bovine ovarian follicular fluid (upper panel) and ovine jugular vein EDTA plasma (lower panel) (S1 Sheffield Method). The sought-for mammalian Candidate 7500 flew in MS from a complex, bacterially contaminated mixture added without treatment to a protein matrix. Calibration (against insulin) varied due to the use of high laser power (operator comment). The mV figures in the header (1.8 = bovine, 13 = ovine) are weak. Strong samples can show up to 1000 mV. The bovine double peak corresponds to an alanine difference, **67mer** vs **66mer** bSgII-70 (S3 Table 2). The ovine double peak corresponds to a lysine and four waters difference, **66mer** vs **65mer**, sSgII-70 (Table 1). All peaks in the lower spectra are predicted to be related to sSgII-70 (S3 Table 1). The item at 4755.85 is a next-integer match to an sSgII-70 predicted **42mer**. The 9509.01 item is double this, suggesting artefactual homodimerization. The lower panel pattern has been reproduced on more purified uncontaminated upstream precipitate (S3 Fig. 5), while the upper panel has also been replicated (S3 Fig. 4). MS courtesy of Carolyn Carr, University of Oxford, Oxford, UK.

**Figure 5.**
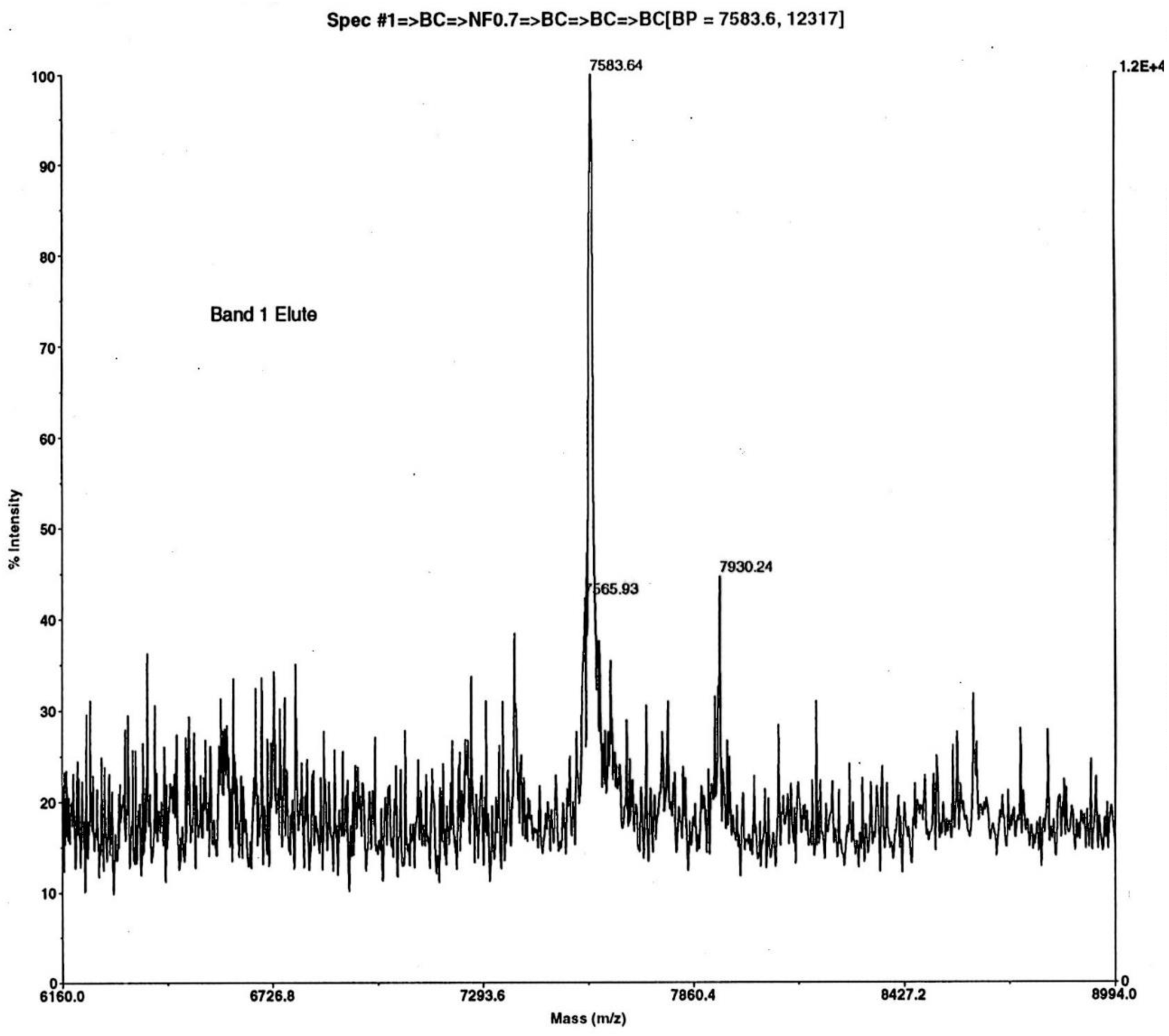
MALDI mass spectrum of a solid ‘upstream precipitate’ that formed on the 3 kDa membrane during a 3-30 kDa ultrafiltration (with sodium azide) of ovine jugular vein EDTA plasma (S1 Sheffield Method) – without the bacterial contamination of Fig. 4. Matrix and calibration as Fig 2. Acquisition mass range 2500-25000 Da. Electroeluted SDS-PAGE Band 1 at ∼7 kDa showed cell-inhibitory bioactivity in vitro. Edman sequencing of Band 1 from a prior (contaminated, azide free) purification provided the 14-residue N-terminal sequence MKPLTGKVKEFNNI (EPL001). This matched anionex Beale 4, the minimal sequence obtained from maximally purified (bacterially uncontaminated) material. Gel electroelution, as used here, provides a cleaner spectrum than solvent extraction (S3 Fig. 5), albeit with the same base peak, an sSgII-70 **65mer**, lacking –KANNI (Table 1).

**Figure 6.**
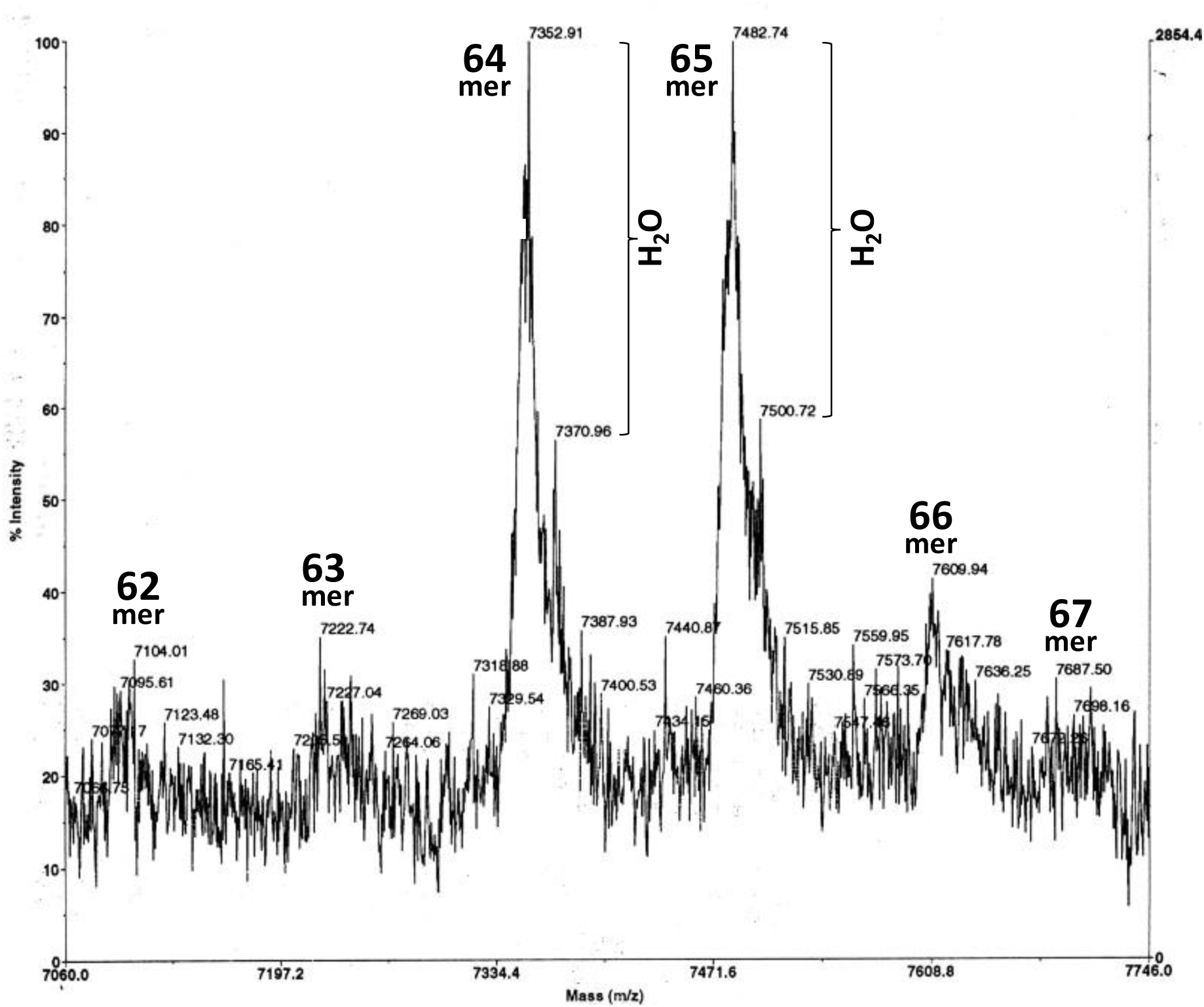
Bovine Candidate 7500. MALDI mass spectrum of bovine ovarian follicular fluid, analysed for variations in bSgII-70 C-terminal truncation and water differences. Anionex Fraction 24 (i.e. late eluting), after spin and gel filtration. The statistical significance of proposed matches are at the >99% confidence level (chi-squared test), as in the sheep (S3). Matrix and calibration as Fig. 2.

**Table 1.**
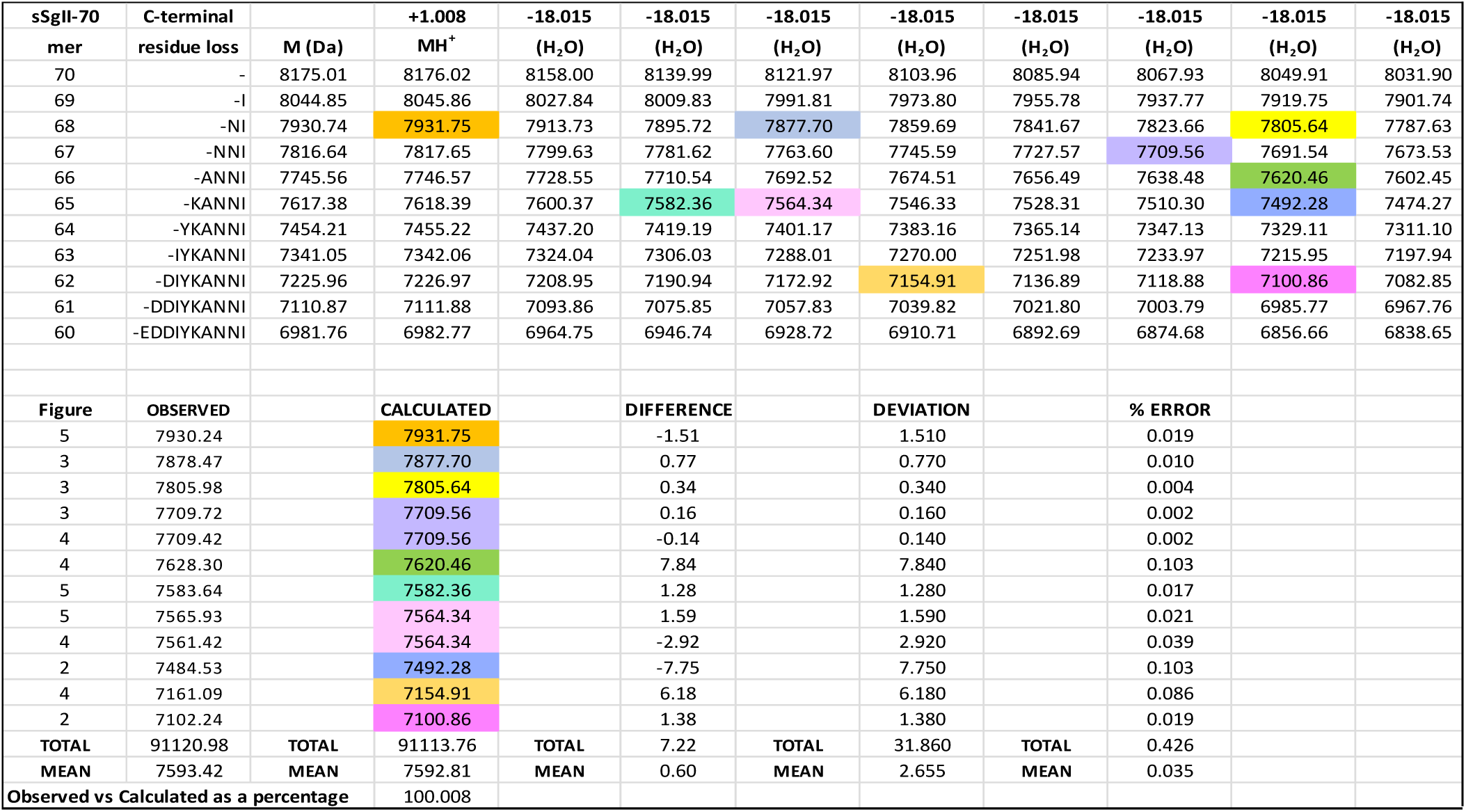
MALDI peak heterogeneity analysis based on C-terminal truncation and water losses for ovine Candidate 7500 identified as sSgII-70. Masses are for the native **70mer** polypeptide sSgII-70 down to a **60mer** (with predictions generated by Expasy Compute pI/Mw and doubled-checked via an MS/MS fragment ion calculator: see S3). A mean deviation of *m/z* **2.66** is delivered at a mean error (i.e. difference as a percentage of match) of **0.035%**; seven integer or next-integer matches (**58**%); ∑Observed (n = 12) 91120/∑Calculated Matches 91113 x100 = **100.01**%; and chi-squared P = 0.024, significant (S3 Table 5).

Counting Candidate 7500 as Candidate I and EPL001 as Candidate XI, there were eight other candidates from the physicochemical purification of sheep material, including fibrinopeptide A and androglobin. Seven of these eight were ruled out on the basis of size, contra-activity and insubstantiality. The eighth, Candidate IV, is ‘Luteal phase entity’ at *m/z* 3450. This came to be interpreted as a fragment of Candidate 7500 (see ‘sSgII-70’, 3000s analysis). Candidate X from the rat, NXPE family member 4 (Q5XI89, formerly FAM55D), was discounted on the basis of predicted pI. Candidate XII will be analysed later under the rubric ‘Likely SgII related’. The dismissal of all but three of the twelve candidates for the inhibitory gonadal influence amounts to the discounting of a series of alternative hypotheses.

Besides the candidates described, there could in principle have been other entities present in the anionex samples undetected by MALDI, though a small molecule hunt using this MS modality detected nothing and neither did one up to *m/z* 25,000. Meanwhile a ‘no preconceptions’ investigation of anionex fractions using ESI MS showed no significant peak large or small, including the frontrunning Candidate 7500 (= Candidate I). Purification reproducibility waned in regard to this candidate, for reasons that only became clear later with the recognition of factor grand fragmentation. Candidate 7500 had not been lost. Bits fall off its C terminus, as evidenced by MALDI spectra. Factor elusiveness vitiated de novo sequencing by MS/MS using trypsin proteolysis (see later in this section) and compromised the use of a polyclonal antiserum (hereafter ‘antibody’) raised against EPL001 (= Candidate XI), when such became available, to aid identification by subtraction from spectra.

Left standing by the foregoing analysis are Candidate I (= **Candidate 7500**), Candidate XI (= **EPL001**) and Candidate XII (= **‘Likely SgII related’**).

### Candidate 7500

Candidate 7500 is not a unitary entity, but a perplexing range of MALDI peaks of *m/z* 7-8000 (Figs. 2-6; S3). In MALDI the sample is applied to a metal target plate with a low molecular weight UV absorbing organic matrix in vast excess to aid desorption and ionization. When this mixture dries it forms crystals, to which a UV laser imparts energy. The matrix heats up rapidly and sublimates, along with the sample molecules co-crystallized with it. The positive ions so produced, most singly charged, ablate into the vacuum and are accelerated in an electrical field into an electrostatic flight tube before being reflected between magnets, for separation on the basis of *m/z* ratio. In a Time-of-Flight (TOF) setup the ions are accelerated to equal energy for mass measurement. MALDI-TOF MS offers high sensitivity, detecting proteins and peptides for example at low femtomole levels. MALDI is fairly tolerant of salts and other background molecules, simplifying sample preparation. It is regarded as a soft ionization method, which aids sensitivity, except that high laser energy had to be used in the present project to visualise exiguous material, contributing to a high noise to signal ratio. Hitting material hard and often with a laser to visualize peaks can cause sample decay and disturb calibration. The weakness of some Candidate 7500 peaks prompts concern that they represent contaminants. One potential cause of heterogeneity in MS is SDS, as used in electrophoresis, but Candidate 7500 showed multiple peaks even when SDS-PAGE was *not* involved (e.g. Fig. 4).

Confusion is dissipated by the realisation that the multiple *m/z* 7-8000 peaks in MALDI probably arise from a single polypeptide subject to molecular losses in the form of C-terminal aa residues and water molecules (‘dual decrementation’). The decryption key is Candidate XII, in the form of ‘Likely SgII related’, with an MS analysis given in full later (see ‘SgII-70’). Let it be said here that while the dually decremented MS data analysis of Table 1 works in pulling observations tightly into a scheme of C-terminal truncation and water losses, it is problematic on the basis of a dearth of consonant literature reports relating to MALDI, an intact mass form of MS.

Candidate 7500 is eluted in anionex by low salt (0.1-0.2 M NaCl) and high salt (0.8 M NaCl). Bovine ovarian follicular fluid was subjected to the standard purification procedure of spin filtration, gel filtration and anionex (S1 Babraham Method). The final step yielded 24 fractions (F1-24), six of which boasted *m/z* 7-8000 moieties: **F10** (7170, 7470), **F11** (7500), **F19** (7062, 7133), **F20** (7065), **F21** (7368, 7498) and **F24** (7352, 7482). The mass spectrum for Fraction 24 is shown as Fig. 6. No pattern is discernible in terms of *m/z* as between low-salt and high-salt forms of Candidate 7500. Concordant with acidity indicated by successful anionex separations was the failure of two laboratories to furnish MS candidates using cation exchange chromatography. Both early and late eluting anionex fractions containing Candidate 7500 reduce the viability of BMCs in vitro, producing a ‘double dip’ histogram, though a single late-stage dip became more characteristic over time. In one purification run involving anionex HPLC, 17 fractions of ovine plasma returned these figures in terms of BMC viability (% control), with Candidate 7500 eluting early and late and with the deepest inhibitions emphasized: 98, **72**, 79, 111, 92, 91, 81, 88, 94, 85, 93, 86, 93, **71**, **73**, 90, 77. The low-salt form of Candidate 7500 provided initial assay data (e.g. Fig. 1) and MS findings, before being lost to view. Disintegration of the early form can be suspected, probably because of unappreciated effects of a change of procedure (S1 Sheffield Method). The tenaciously bound high-salt anionex form of Candidate 7500 became the focus, providing inhibitory exemplifications in vitro (S2) relating for example to human breast and prostate tumour cell viability, rat cardiac hypertrophy, vascular smooth muscle growth (rat aortic stems cell numbers halved over 72h, P<0.05, Bonferroni t-test) and angiogenesis. In the last-mentioned application, pooled anionex Fractions 28 & 29 of bovine ovarian follicular fluid containing Candidate 7500 inhibited venule formation in human umbilical vascular endothelial cells to a comparable degree (>50%, ANOVA, significant: S2) to the well-known potent anti-angiogenic compound suramin administered at 20µM.

When a late-eluting anionex fraction containing Candidate 7500 was subject to C18 reverse phase chromatography (RP-HPLC) a bimodal distribution occurred, as in the prior anionex, in regard to both the presence of Candidate 7500 and BMC inhibition in vitro. These bimodalisms indicate that Candidate 7500 is amphipathic. Ovine plasma anionex late-eluting Fraction 41 had MALDI peaks at *m/z* 2375>6967>7628. In 17 reverse phase fractions, peaks were seen only in fractions 3-7 and 11-16. In the first wave the peaks were 5400>7626>2251, in the second 5400>5800>7630>2255 (S3 Fig. 3). Disintegrations of Candidate 7500 were deemed to have occurred, providing fragments that were themselves bimodally distributed. A separate study involved evaluation in two laboratories of the same anionex sample. The focus was on ovine follicular fluid anionex late-eluting Fraction 29. This was subject to MALDI (matrix sinapinic acid, calibration against carbonic anhydrase, as for the reverse phase analysis) in the East of England at Babraham. There was a main peak at 7095, medium peaks at 5454 and 4273 and a small peak at 7896. Fraction 29 was then analysed shortly afterwards in Central England at Harwell Laboratory, again using MALDI (insulin calibration). There was a main peak at 4258, with subsidiary peaks at 4387, 4306, 2946 and 2479. Peaks in the 7000s were absent. This supports the interpretation that Candidate 7500 is labile (i.e. disintegration prone, unstable) and that major subsidiary peaks represent ‘Grand Fragments’ of Candidate 7500, a view that will be developed fully later on (see ‘sSgII-70’).

A uterus-only version of the rat organometric assay in vivo was carried out on a 3-30 kDa fraction of sheep ultrafiltered plasma subject to anionex HPLC (S1 Sheffield Method). The zero-baseline uterine weight was provided by rats injected intraperitoneally with PBS for four days. Three groups of fractions lacking Candidate 7500 (validated by MALDI) were tested: pooled anionex Fractions 1-11 were associated with mean relative uterine weights *below* baseline by 7%; Fractions 12-22 were *above* baseline by 6%; and Fractions 25-32 were also *above*, by 8%. None of these findings was significant. Fractions 23 & 24, unlike the others, contained (late-eluting) MS Candidate 7500. Pooled, these fractions were associated at termination with a uterine weight 22% *below* that of controls (P<0.002, significant, ANOVA).

Candidate 7500 material almost halved inhibition of compensatory renal growth in the unilaterally nephrectomised male rat (S2). Late running anionex HPLC fractions were used, containing and lacking Candidate 7500 (validated by MALDI). As much Candidate 7500 was used as was present in 360ml of ovine pooled ovarian follicular fluid. Administration was via an osmotic mini-pump implanted into the residual kidney’s pelvic region. Values for the residual kidney (i.e. post-treatment) are expressed in terms of the removed kidney (i.e. pretreatment) as % change, mean +/–SEM (t-test, equal variance, all findings significant except as indicated), control versus test: kidney wet weight, 41.7 +/– 2.7 vs 23.2 +/– 2.6 (P<0.005); dry weight, 25.9 +/– 5.9 vs 13.5 +/– 1.1 (P<0.012); protein, 65 +/– 49 vs 28 +/– 14 (P<0.2, ns); and DNA, 81 +/– 23 vs 42 +/– 15 (P<0.05).

Sheep plasma in one series was subject to the standard purification procedure of spin and gel filtration followed by anionex (S1 Babraham Method). In six separate fractionations involving ovarian venous plasma Candidate 7500 was seen in MALDI. In contrast Candidate 7500 was *not* seen in jugular vein plasma from two OVX ewes, supporting the factor’s ovarian provenance. In the search for an inhibitory *gonadal* influence, then, Candidate 7500 is fittingly ‘ovary plus’ and ‘ovariectomy minus’: OV+/OVX–. Candidate 7500 was present in the blood of hypophysectomised ewes one day and seven days post-surgery.

Candidate 7500 is higher in the follicular phase of oestrus, suggesting an appropriate reproductive relatedness (S2). Ovine blood serum was collected at different phases of the oestrous cycle and purified in the usual way to anionex HPLC fractions. There was a peak of Candidate 7500 in the follicular phase with a second smaller peak in late luteal. In MALDI the combined counts for Candidate 7500 in late-eluting fractions 19-21 were: follicular, ∼13,140; early luteal, ∼1770; late luteal, ∼4570. Inhibitory activity in the BMC viability assay in vitro is stronger in sheep blood from the follicular phase than that from the luteal phase. Ovine samples collected in Spring were higher in Candidate 7500 and more active in vitro than were samples collected in Summer or Winter.

During an upscaled 3-30 kDa ultrafiltration purification, a precipitate formed on the 3 kDa filter membrane (S1 Sheffield Method). The supernatant, designated Sheep Plasma Filtrate, was inhibitory in the BMC assay, as was the solid precipitate – dubbed ‘upstream precipitate’– mixed together with cell medium. MALDI of whole upstream precipitate startled by providing Candidate 7500 (Fig. 4). Washing the solid caused the loss of Candidate 7500 (S1 Fig. 11). An aqueous extract of upstream precipitate was actively inhibitory in vitro. The bioactivity thus followed the whereabouts of Candidate 7500, with an aqueous extract of the upstream precipitate providing the Edman sequence MKPLT/GKVKxFNNIK/IGF/Y/DxF/VI/VI, which is denoted the ‘EPL001 Extension’ (S1 Table 1).

Modelling of bioactivity in vitro saw Candidate 7500 anionex fractions inhibit cardiac hypertrophy by >20%, compared with fractions lacking Candidate 7500 (Hart, 2008). Measured in cultured myocytes from neonatal rats was the RNA ratio of atrial natriuretic factor, a marker of hypertrophy, to glyceraldehyde 3-phosphate dehydrogenase, a housekeeping enzyme, at the end of a 24h exposure period. Microscopic analysis showed no extra cell death in the Candidate 7500 group but suggested a reduction in cell size. In other exemplifications in vitro there were fewer smaller cells, with reduced cell division accompanied by cell death due to physiology (apoptosis) rather than toxicology (necrosis) (Hart, 2000). One interpretation of these data is that the ovary elaborates into the circulation a cell shrinking agent, Candidate 7500.

Cell fate has been assessed in the rat bone marrow cell assay by various methods (Hart et al 2017), including cytometry using an Annexin-V-Fluos Staining Kit (Sigma-Aldrich, Dorset, UK). Among cells exposed to sheep test plasma anionex fractions (i.e. containing Candidate 7500) pre-apoptotic and apoptotic cells comprised about 60% of the population, compared with 3-4% in controls, with both healthy and necrotic cells less prevalent in the test population. A dose-response curve for apoptosis has been obtained. This was in regard to the hypothalamic aspect of the inhibitory hormonal proposition (Hart, 2014). An anion exchange fraction of aqueous extract of rat hypothalamus was evaluated via image analysis for proliferative and apoptotic action on a mixed population of rat bone marrow cells in vitro (Hart et al, 2017). The late Candidate 7500 fraction was used, eluting at 0.8 M NaCl. Control cell numbers increased in numbers by more than 40% in a 24h period. Test cells receiving a single dose of the anionex fraction maintained a steady population. Among test cells exposed to a double dose of fraction death began to occur after 5h, with 90% of the cells having undergone apoptosis by 24h, as judged morphologically. A triple dose of fraction caused apoptosis to begin after 90 minutes. In under 4h all the cells were dead (personal communication, Ben Ferneyhough, Systems Biology Laboratory UK, Abingdon, Oxfordshire, UK). As well as being observed in primary cultures of rat bone marrow cells, apoptosis has been induced by Candidate 7500 material in a mouse bone marrow cell line (SR4987) and in a variety of human tumour cells lines (e.g. MG-63, MCF7, PC3, DU145) (Hart, 2008). Candidate 7500 preparations tend to deliver stoichiometric (linear) dose-response curves rather than hormone-style sigmoidal shapes (S2).

Attempts were made to elucidate the aa sequence of Candidate 7500 via digestion with trypsin (porcine), which cleaves selectively on the C-terminal side of lysine and arginine residues, together with MS analysis and interrogation of peptide mass fingerprint databases. No convincing hits were seen in a campaign focussed on Candidate 7500 anionex fractions from 13 purification runs, three species (sheep, cow, pig), two source materials (ovarian follicular fluid and blood plasma), multiple MS modalities (MALDI-TOF, Delayed Extraction-MALDI-TOF, QTOF, LCQ Deca XP, ESI-QUAD-TOF and LC-MS/MS) and different online search tools such as Mascot and MS-Fit. A carboxypeptidase was deployed on ovine ovarian follicular fluid fractions to achieve C-terminal truncation in a MALDI study, without productive outcome. A MALDI mass spectrum of a tryptic digest is shown as Fig. 7. This looks unpromising. It is the product of intense lasering: 200 shots/spectrum, as against 100 for Figs. 2 & 3 and 50 for Fig. 5, the other ‘Babraham Ovines’. There are excessive matrix peaks on the left (*m/z* 800s) consistent with analyte in low abundance, plus dominant obvious contaminants. Yet this spectrum yields matches within a bespoke manual analysis (see ‘sSgII-70’).

**Figure 7.**
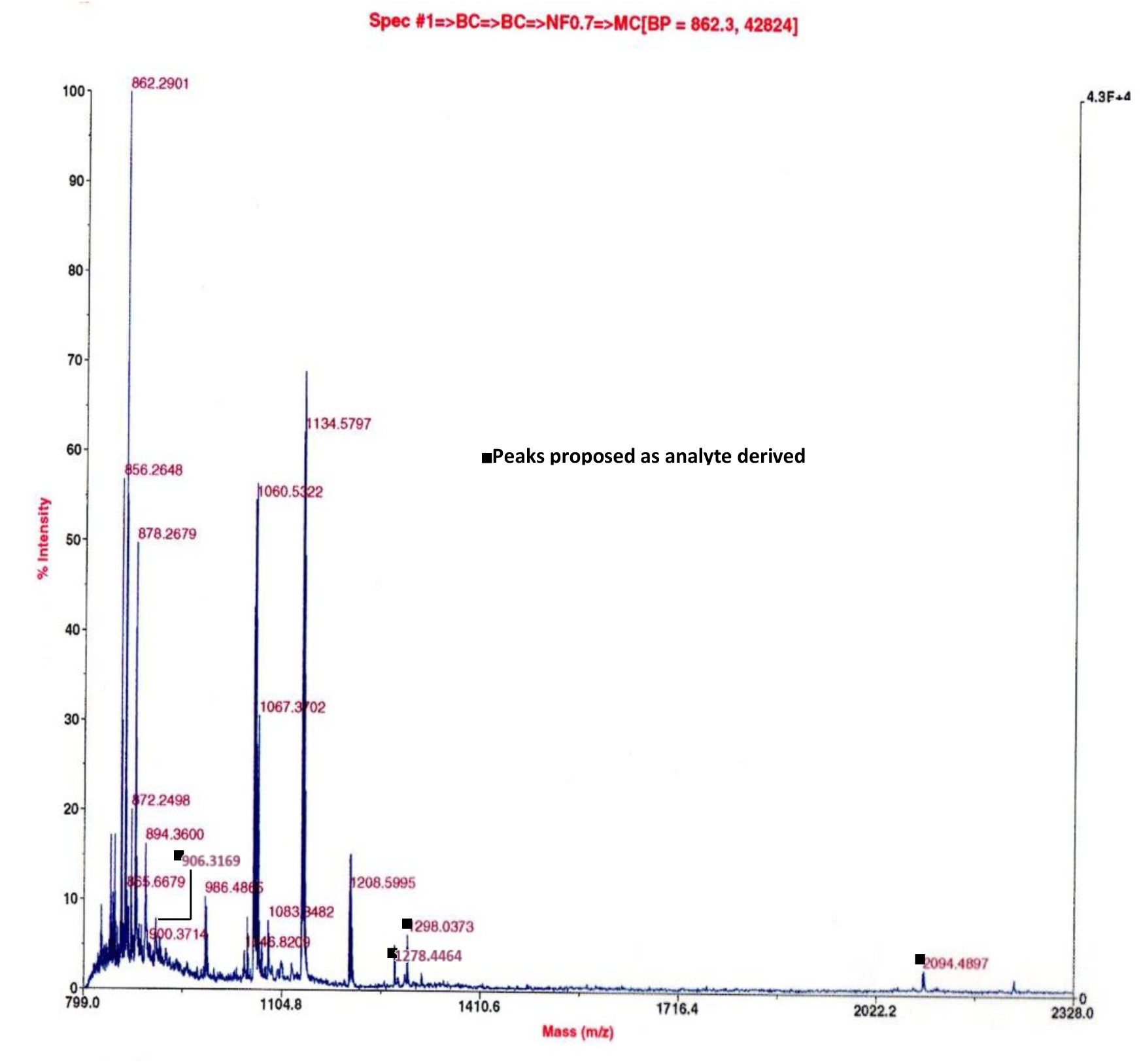
sSgII-70 glimpsed in a MALDI mass spectrum involving ovine ovarian follicular fluid, anionex Fraction 28 containing Candidate 7500, digested with porcine trypsin. Positive ion, reflectron mode, delayed extraction, CHCA matrix, high laser power. Prominent are matrix cluster peaks in the *m/z* 800s and other irrelevancies (e.g. keratin at 1060 & 1134: Mascot Contaminants). Predicted fragments from a tryptic digestion in silico of sSgII-70 find minor peak matches, as per black blocks: (i) Observed *m/z* **906.3169**, Predicted 906.0693, Difference +0.2476 (0.03% above Predicted), sequence (_3_K)TGEKPVFK(R_12_), no missed cleavage; (ii) Observed **1278.4464**, Predicted 1278.5954, Difference −0.1490 (0.01% below Predicted), _1_MLKTGEKPVFK(R_12_), one missed cleavage; (iii) Observed **1298.0373**, Predicted 1294.4123, Difference +3.6250 (0.28% above Predicted), being either (_38_K)LTGPNNQKHER(A_50_), as the Predicted, or the MSO version of 1278 (at 1294.5948), both one missed cleavage; and (iv) Observed **2094.4897**, Predicted 2091.1458, Difference +3.3439 (0.16% above Predicted), (_49_R)ADEEQKLYTDDEDDIYK(A_67_), one missed cleavage. Items (i) and (ii) relate to a proposed reverse splice 9+61 junction at _9_VF_10_. A fifth match is moot, as being at higher (but still low) intensity within the matrix ion group: Observed **872.2498**, Predicted 871.9682, Difference +0.2816 (0.03% above Predicted), (_38_K)LTGPNNQK(H_47_), no missed cleavage. (S3, Digest 5.)

The effect of proteolysis was investigated further when sheep serum was ultrafiltered and subjected to anionex HPLC (S1 Sheffield Method). Control fractions containing no Candidate 7500 provided a baseline viability level in the BMC assay in vitro over 72h, with samples in triplicate. Viability was unchanged when these fractions were exposed to porcine trypsin.

Fractions containing Candidate 7500 were associated with mean cell numbers of 18.5% of the control value (i.e. 81.5% reduction in viability). Trypsinisation of these fractions resulted in cell numbers of 42.2% (i.e. 57.8% reduction in viability). Inhibitory bioactivity was thus reduced by trypsin, albeit incompletely, indicating that Candidate 7500 is proteinaceous but well defended against enzymatic attack. (Similarly heat treatment, 56°C for 1h, only caused partial deactivation, reducing cell death from 88.6% to 54.5%. Charcoal likewise partially reduces activity in vitro.) Ovine plasma still showed antiorganotrophic activity in the rat multi-tissue assay in vivo even after exposure between gel filtration and anionex to agarose-bound bacterial protease (P8790, Sigma-Aldrich, Dorset, UK). Resistance to proteolysis (and heat) is a notable feature of Candidate 7500.

Bioassay-guided fractionation provided proteinaceous Candidate 7500, for which there are multiple incidence-bioactivity correlations, notably OV+/OVX–. The totality of the evidence marshalled here indicates that Candidate 7500 accounts for the tissue-mass inhibitory reproductive hormonal influence. But what is Candidate 7500?

### EPL001

Candidate 7500 is a MALDI candidate from the physicochemical purification but also an SDS-PAGE candidate, being ostensibly responsible for providing a gel band at c7.5 kDa (S1). From this band was obtained ovine aa data via chemical sequencing.

The canonical N-terminal aa sequence MKPLTGKVKEFNNI was designated EPL001, which proprietary designation also refers to a synthetic peptide of the same sequence, a 14mer indeed, where ‘mer’ denotes aa residues. Derived by automated Edman degradation (Applied Biosystems Procise, Foster City, CA, US: referenced as ‘Applied Biosystems Procise’), the EPL001 sequence was seen in whole or in part seven times in separate purification runs involving sheep material that was biologically active (i.e. antiorganotrophic in vivo and antiproliferative in vitro) and contained MALDI Candidate 7500 (S1 Table 1). Notable landmarks are an initial methionine (M), a position-3 proline (P), a mid-sequence valine (V) and a position-11 phenylalanine (F). Five of the seven sequences obtained have enough in common to support a mathematically validated association (Hart et al, 2022). Ostensibly meaningful, EPL001 yet languishes in profound bioinformatic obscurity, garnering no compelling hits in innumerable BLAST searches (Hart et al, 2017). Chemical sequencing (Edman, 1950) was supposed to disclose the character of Candidate 7500, but instead provided a conundrum in its own right.

Edman sequencing and MALDI at Babraham were subjected to an internal calibration demonstrating accuracy by virtue of a contaminant, fibrinopeptide A (S1’s Candidate VI, discounted through contra-activity). An Edman reading, with no subsidiary findings, matched the expected database sequence to the extent of 13 of 14 N-terminal residues, while a MALDI peak from the same anionex fraction at *m/z* 1847 provided a match to a database prediction for ovine fibrinopeptide A of 1848 Da, protonated as MH^+^ to 1849 (S3 Fig. 14b).

EPL001 has proved resistant to molecular biology approaches involving the use of oligonucleotide probes and RT-PCR to find matching sequences in DNA and RNA libraries (e.g. sheep ovary and sheep hypothalamus) and the use of anti-EPL001 antibodies (see next section) to identify cDNA synthesized proteins.

Evidence relating to (ovine) EPL001 is absent from the tryptic digest campaign, which involved trypsin fragmentation and MALDI analysis (S4). Eleven of the thirteen tryptic digests carried out involved purified feedstock that was relevantly ovine. A digest with porcine trypsin in silico (Expasy PeptideMass) of EPL001 (MKPLTGKVKEFNNI), at up to one missed cleavage, yields seven predicted fragments (S4 Table 4). In the MALDI ovine dataset there are no <u>integer</u> matches (i.e. unrounded whole numbers ignoring fractional masses in the form of decimals) to any of these and there are no <u>next-integer</u> hits either (i.e. next unrounded whole number up or down ignoring fractional masses in the form of decimals), except to a predicted fragment at 863 Da (VKEFNNI). This latter match is in the form of overly prominent peaks at *m/z* 861/2 in nine of the eleven trypsinisations, being a known cluster of the matrix chemical CHCA, i.e. α-cyano-4-hydoxycinnamic acid. (This peak is also present in OVX material lacking Candidate 7500, supporting a claim of irrelevance.) Notable is an absence of <u>integer</u> or <u>next-integer</u> matches to the one predicted EPL001 fragment comfortably outside the matrix region and which does not match the mass of any known contaminant: MKPLTGKVK (1002 Da at one missed cleavage, with the MSO variant at 1018). Concluding, there is a lack in samples of predicted tryptic fragments of EPL001.

The unproductivity of bioinformatics, molecular biology and tryptic digestion lends three-way support for the conclusion that EPL001 does not represent a true-to-life N-terminal sequence.

EPL001 has been investigated in terms of bioactivity as a synthetic 14mer peptide to which antibodies have been raised. The molecular formula of EPL001 is C_73_H_129_N_19_O_20_S_1_, while the molecular weight is 1618.95, with a theoretical pI of 9.70 (Expasy Compute pI/Mw). As a synthetic peptide EPL001 surprised by being anti-organotrophic in its own right, dose-dependently, in an assay in vivo of rat compensatory renal growth after unilateral nephrectomy (Haylor et al, 2009) and also by modulating reproduction in the nematode *Caenorhabditis elegans* (Davies & Hart, 2008). Both these activities are aspects of the inhibitory hormone concept (Hart, 2014), with the inhibition of compensatory renal growth seen with EPL001 also having been seen with Candidate 7500, as described earlier. Crucial in compensatory renal growth is the endogenous mitogen IGF-1, insulin-like growth factor 1 (P05019; UniProt numbering throughout) (Haylor et al, 2000). IGF’s effects are evidently counteracted by EPL001 and Candidate 7500, while being potentiated by an anti-EPL001 antibody (see later). Referring to EPL001, the likelihood is low of a project-unrelated peptide having both the predicted activities of antiorganotrophism (tissue shrinkage) and reproductive modulation. Additionally, EPL001 enhanced lifespan in the nematode (Davies & Hart, 2008).

EPL001 has been shown to be inhibitory in multiple cell assays in vitro. The peptide reduced cell division for example in human prostate cancer cells in cultures, as judged by the incorporation of tritiated thymidine, and reduced the proliferation of rat BMCs (Hart et al, 2022). Human marrow derived mesenchymal stem cells in culture exposed for 14 days to EPL001 tethered to cover slips were present in ∼25% lower numbers than controls (op. cit.). Analysis using fluorescence-assisted cell sorting (FACS) indicated a lower proportion of cells in the later stages of the cell cycle, with 26.95% of control cells positive for Ki67 (nuclear cell cycle marker absent from G_0_) versus 18.92% of tests. A lower average cell size is implied by these results. EPL001 thus provides fewer smaller cells, as do ovine anionex fractions containing Candidate 7500.

Intracerebroventricular infusion of EPL001 in sheep was associated with elevated growth hormone in peripheral blood and reduced prolactin (Hart et al, 2022). The 14mer EPL001 peptide thus shows a repertoire of relevant bioactivities while presumptively only representing 20% of the sought-for proteinaceous entity of ∼70 aa.

### Anti-EPL001 antibodies

The raising of antibodies to the 14mer synthetic peptide EPL001 has been described previously (Haylor et al, 2009), including in regard to what became the lead antibody, a goat polyclonal antibody designated G530 (Hart et al, 2017; Howlett et al, 2019). In western blots this latter antibody provided bands in early and late eluting anionex fractions of serum from ovary-intact sheep, in the same incidence pattern as Candidate 7500 (S1 Fig. 4). Candidate 7500 is OV+/OVX–; likewise the circulating endogenous antigen of an anti-EPL001 rabbit polyclonal antibody, which antigen in OVX– mode was lacking from 3-30 kDa blood plasma ultrafiltrate of OVX sheep (Hart, 2008). These correlations support a connection between EPL001 and Candidate 7500.

Western blots using the G530 anti-EPL001 antibody have visualized Candidate 7500 bands in a range of biological materials, including sheep plasma, rat PC12 conditioned media and rat hypothalamic aqueous extract (Hart et al, 2017). The synthetic antigen EPL001 is MKPLTGKVKEFNNI. Dot blotting with EPL001 ‘Front 6’ and ‘Back 8’ peptides has indicated that the epitope (singular for parsimony) within EPL001 recognised by the anti-EPL001 goat polyclonal resides exclusively in the Back 8: K_7_V_8_K_9_E_10_F_11_N_12_N_13_I_14_. Such an epitope is also what is being seen by the antibody endogenously, as confirmed by preabsorption of ∼7 kDa Western bands of rat hypothalamus aqueous extract with the Back 8 peptide but *not* with the Front 6, M_1_K_2_P_3_L_4_T_5_G_6_ (Hart et al, 2017). The evidence from IHC peptide preabsorption studies indicates that the epitope within EPL001 of the anti-EPL001 G530 antibody is **<u>KEFNNI</u>** (Howlett et al, 2019). The epitope within the antibody’s mammalian endogenous antigen has been deduced to be a discontinuous version of this: **<u>KE·F·NNI</u>**. A 14×14 sequence grid analysing EPL001 has been described (Hart et al, 2022) and will be introduced fully in the next section (‘Likely SgII related’). The residues of the endogenous epitope provide the plot’s southern fringe, cross-validating grid and epitope.

Immunohistochemistry (IHC) images of high specificity and endocrine relevance have been provided by anti-EPL001 antibodies (Hart et al, 2017). Hypothalamic staining in the rat was evident in the arcuate nucleus, with individual neurons staining also in the retrochiasmatic nucleus. Staining was apparent in individual neurons in the ovine lateral and ventromedial hypothalamus and preoptic area, with heavy staining in the palisade (neuroendocrine) region of the median eminence, with axonal beading betokening transport. Staining was not seen in either the sheep or rat pituitary. Staining was evident in human testicular germinal epithelium (also spermatogonia), without appearing in the testicular interstitial Leydig cells, whence testosterone. The rat ovary showed moderate staining in theca, granulosa, follicle cells and follicular fluid. Discrete *neuroendocrine* cells stained in intestinal and other human organs – an interpretation supported by serial-section co-localisation using the neuroendocrine marker chromogranin A. IHC findings were thus consistent with a sought-for hormonal factor. Staining of neuroendocrine relevance was also seen in embryos of the fruit fly (*Drosophila melanogaster*).

Consonant with the IHC findings, ovine median eminences were used at one stage in the purification campaign (Hart et al, 2022). An acid extract was subjected to anionex and RP-HPLC. Immunoreactivity was evinced by a single fraction using an anti-EPL001 goat antibody. Even so, no data were obtained from Edman sequencing or MS/MS. The endogenous antigen to the anti-EPL001 antibody thus retained an inscrutable elusiveness yet the activity of said endogenous antigen was demonstrable in sundry systems using the antibodies. For example, rat bone marrow cell controls in an assay in vitro (method in Hart et al, 2017, with 2D video image analysis using ImageJ software) increased in individual size by an average of about 11% over 24h. Test cells exposed to a 23% solution of unpurified goat polyclonal anti-EPL001 G530 antibody grew by 29% (personal communication, David Howlett, coauthor). This finding is consistent with the immunoneutralization of a cell-size-limiting antigen, presumably present in the medium. Administration of an anti-EPL001 antibody caused compensatory renal growth to overshoot in a rat assay involving unilateral nephrectomy, also consistent with the immunoneutralisation (IN) of an endogenous inhibitor (Haylor et al, 2009).

A prototype ELISA (enzyme-linked immunosorbent assay) has been developed (S2) based on an anti-EPL001 rabbit polyclonal antibody (Hart et al, 2017). Human males had 2-3 times as much of the analyte in plasma as did human females, a result that awaits replication.

Anti-EPL001 polyclonal antibodies have been deployed in immunoaffinity column purifications to identify the endogenous antigen without decisive outcome (Hart et al, 2022). The identification impasse was ultimately breached by these means: (a) antigen capture by immunoprecipitation (IP, with LC-MS) rather than immunoaffinity columns; (b) superseding automated Edman degradation with Orbitrap MS sequencing after trypsinisation; (c) consideration of a broader MW range rather than for example 3-30 or sub-10 kDa material, taking in anything that is water soluble; and (d) use of biological starting materials preserved via formalin fixation crosslinking, with antigen retrieval, since this approach worked in IHC.

### ‘Likely SgII related’

The sought-for antiorganotrophic hormonal influence is deemed ‘Likely secretogranin II related’. This is a triangulated result by virtue of IP/LC-MS purification, IHC and bespoke bioinformatics. SgII relatedness is the key piece in the factor identification puzzle (Fig. 8). SgII is a neuroendocrine protein of the granin family that regulates the biogenesis of secretory granules and is a prohormone.

**Figure 8.**
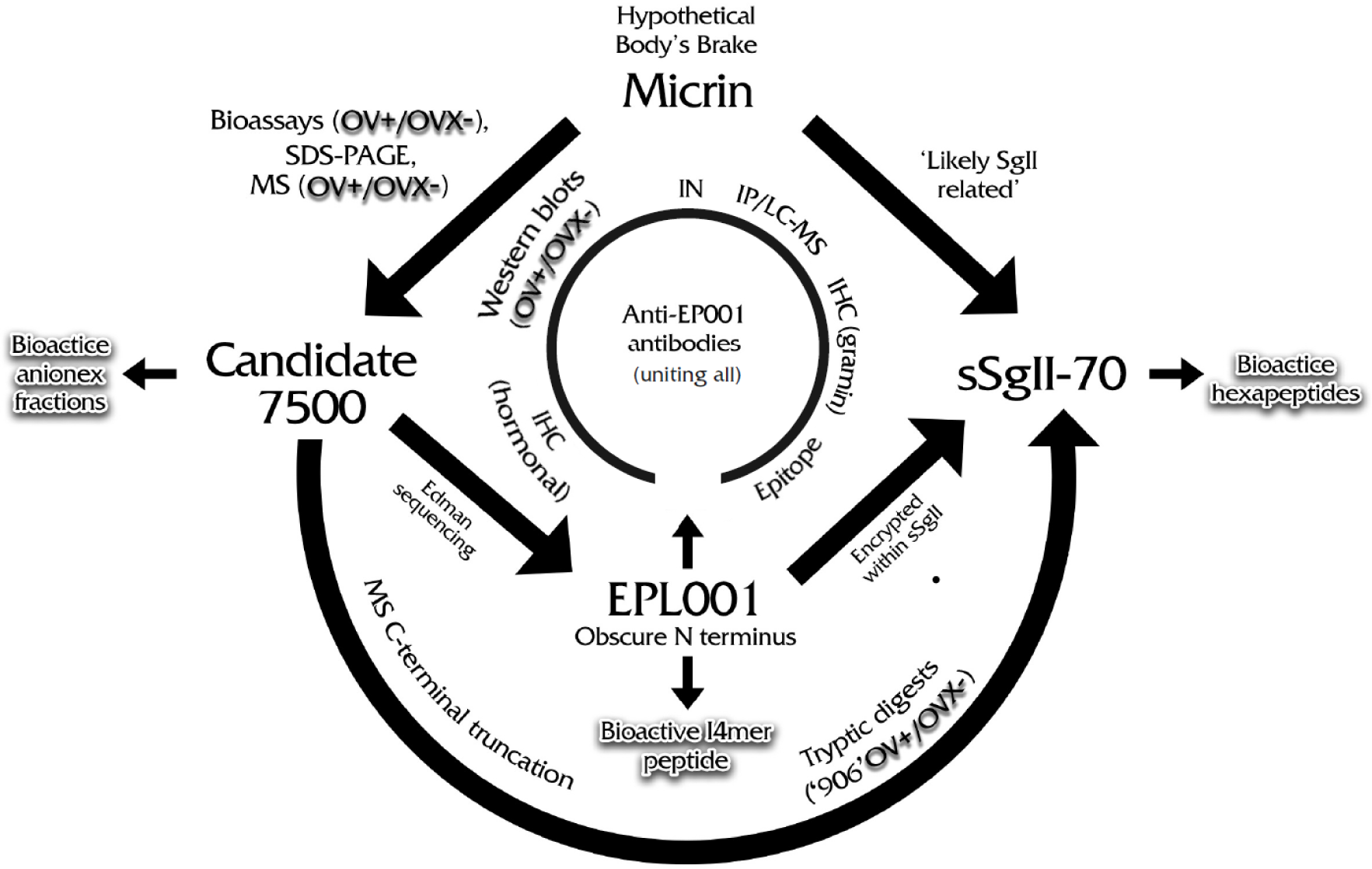
Puzzle. IN = immunoneutralization, IP/LC-MS = immunoprecipitation/liquid chromatography-mass spectrometry, IHC = immunohistochemistry, OV+/OVX– = ovary intact material presenting positively/ovariectomised material presenting negatively.

‘SgII relatedness’ arose on the basis of the G530 antibody’s use in an IP/LC-MS antigen-capture campaign directed at an aqueous extract of rat hypothalamus (Hart et al, 2017). The rat hypothalamus is IHC+ and a neat aqueous extract is antiproliferative and proapoptotic in an assay in vitro involving rat BMCs, with this inhibition subject to prior immunodepletion by the anti-EPL001 antibody. For antigen capture the same antibody was bound covalently to resin beads, with control preparations preabsorbed with EPL001. Cryopreserved material (i.e. formalin *unfixed*) delivered no credible candidate. More productive was formalin fixation with antigen retrieval, simulating what had worked in IHC. Presumed factor lability had apparently been countered. A sole candidate emerged from two antigen retrieval approaches, RIPA buffer and citrate buffer. This was identified by the MS software as ‘SgIIvar’, Q8CGL8, a half-sized splice variant of rSgII, P10362 (Gene: *SCG2*). On the basis of preabsorption evidence from Westerns this has been interpreted as an intermediate in the intracellular processing towards secretion of an SgII proteoform, where ‘proteoform’ refers to all protein variants of a single gene including post-translational modifications and aa sequence variants. A parallel IP/LC-MS exercise involved embryos of *D. melanogaster*. Brought forth from formalin fixed and antigen retrieved material was the uncharacterised protein Q9W2X8 (Gene: *CG12644*), potentially explaining neuroendocrine staining in this species. The rSgII intermediate proteoform and fly Q9W2X8 display the same partial epitope (NNI) and a version of the rat protein’s MS ID peptide is present in the fly protein. A version of Q9W2X8’s N-terminally located MS ID peptide is lacking from the N-terminally truncated 322-residue rSgII proteoform but is present in the 619-residue full-length rSgII, in the same relative position. Fly Q9W2X8 shows detailed SgII-relevant sequence homology, notably in regard to sorting domains and dibasic residues. SgII is a member of the granin family of secretory-granule proteins, which includes chromogranins A and B (CgA & CgB), SgIII and several other granin-like proteins (Bartolomucci et al, 2011). These entities are involved in the biogenesis and operation of intracellular secretory vesicles as part of the regulated secretory pathway by which neuropeptides and protein hormones are provided for export from endocrine, neuroendocrine and neural cells. Granins are soluble, acidic, heat-stable, calcium-binding proteins that are prohormones in their own right, giving rise to (stoichiometric) bioactive peptides (Zhao et al, 2009). SgII is encoded by gene *SCG2*, which in the rat is expressed in the brain, adrenals and testes, as judged by RNA sequencing, without significant expression elsewhere (NCBI Gene ID 24765). The protein is tyrosine-sulphated by way of post-translational modification. SgII exhibits no sequence homologies with CgA & CgB and is poorly conserved compared with them. Only short peptides known as secretoneurin (SN) and EM66, proteolytically processed from the central core region of SgII, show a high sequence similarity among the vertebrates (Zhao et al, 2009). SN’s activity profile includes immune and inflammatory modulation, while EM66 may be involved in modulating food intake.

Hypothalamic IHC staining is suggestively similar for an anti-SN antibody (Marksteiner et al, 1993), an anti-EM66 antibody (Boutahricht et al, 2007) and anti-EPL001 antibodies (Hart et al, 2017), notably in regard to a plexus of stained neurons in the median eminence. EM66 visualizes additionally in the rat pituitary (Montero-Hadjadje et al, 2003), while the antigen of the anti-EPL001 antibody does not visualize in the sheep or rat pituitary (Hart et al, 2017). Neither peptide can be the factor sought in the present enquiry, as SN & EM66 both lack the full complement of EPL001 residues and neither can in principle be seen (in terms of epitope possession) by the main anti-EPL001 G530 antibody (Howlett et al, 2019).

Granted EPL001’s ovine provenance, a bioinformatic analysis has been presented based on that part of the TrEMBL database devoted to *Ovis aries* (Howlett et al, 2019). The evidence from IHC peptide preabsorption studies is consistent with the mammalian epitope of the G530 antibody used in the IP/LC-MS purification being KE·F·NNI. Of 26,443 ovine predicted proteins 1,100 proteins contain one or more NNIs. These 1,100 proteins comprise the ovine ‘NNI-ome’, with 1,181 NNIs in all. The assumption is that one of these motifs relates uniquely to the NNI in EPL001’s sequence MKPLTGKVKEF**<u>NNI</u>**. Within the NNI-ome’s 1,100 proteins there are 22,207 methionine residues, discounting signal sequence initiator Ms. This is the pool of candidates for the M at the start of EPL001. Ignore NNI and focus on EPL001’s other 11 residues. The Method of Exclusion can be used to eliminate all candidate Mxxxxxxxxxx sequences bearing residues *not* present in EPL001, i.e. ARNDCQHISWY, plus M, as that is used up as the first residue. Of Mxxxxxxxxxx sequences in the ovine NNI-ome 38 are composed exclusively of the 9 unique residues in EPL001-less-NNI, counting repeat aa as one. None has all 9. Five have 8, the rest having from 7 down to three. The five with 8 EPL001-less-NNI residues are as follows, described by similarity with other mammalian proteins: W5Q754 (titin, structural, MLKKTPVLKKG); W5PWS5 (dynein, structural, MLFVGPTGTGK); W5PP00 (ubinuclein-2, nuclear, MPKVVPTLPEG); W5PWP9 (hormonally up-regulated neu tumour-associated kinase, enzyme, MLTGTLPFTVE) and W5QEU8 (secretogranin II, secretory vesicle protein, MLKTGEKPVEP). The stand-out candidate from amongst these five is the last, SgII, an established prohormone, the match being to SgII’s second sorting domain.

Secretogranin II has two main sorting domains that can act independently to direct the protein within the cell’s cytoplasm into secretory granules of the regulated secretory pathway (Courel et al, 2008). In the second main sorting domain of sheep SgII there is a curiosity, as is evident from the NNI-ome bioinformatics: a sequence of aa (‘sSgII-9’) that is a 9/14 shuffled version of EPL001, with the latter as _1_**<u>M</u>**KPLTG**<u>K</u>**VKEFNNI_14_ and sSgII-9 as _367_ **<u>M</u>**LKTGE**<u>K</u>**PV_375_ (Fig. 9). Just the two residues emphasized are in register.

**Figure 9.**
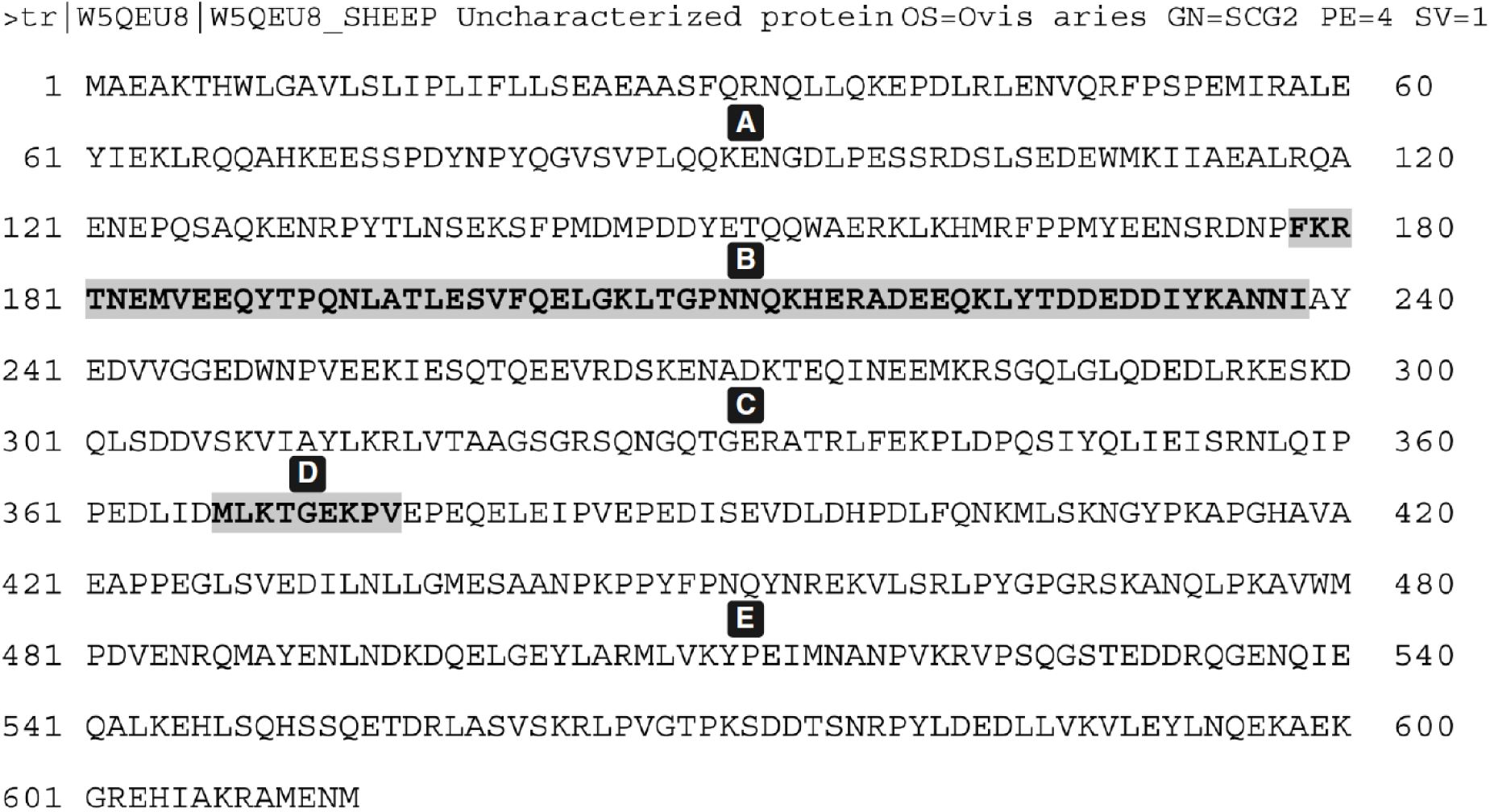
Sheep secretogranin II preprohormone (UniProt W5QEU8, FASTA format). Includes sSgII signal sequence (M1-F30; Courel et al, 2008), which is the ‘pre’ in ‘preprohormone’. sSgII is postulated to give rise to a secreted proteoform, sSgII-70, the product of a 9+61 reverse peptide splicing (DB in an A-E modularisation, with residues emphasized). Yet SgII is self-aggregating, admitting the possibility that the 9mer D and 61mer B units of sSgII-70 derive from different SgII molecules.

Missing for a full ovine match with EPL001 are residues K, F, N, N, I. The last three can be accounted for by the partial epitope _236_NNI_238_. This commands attention, especially since this part of sSgII, with polyanionic groups separated by hydrophobic clusters, can be suspected itself of having sorting relevance (Hart et al, 2017). Now with a 12/14 anagram – where an anagram is defined as a word or phrase formed by rearranging the letters of another word or phrase – what of the missing K and F? Among the 43 Ks and 10 Fs in sSgII there is only one pairing: _178_FK_179_ (Fig. 9). Taking this doubleton for the sake of argument as the provider of the absent residues the contribution of ‘sSgII-14’ to this admittedly bizarre 14/14 ‘match’ is as follows: _367_MLKTGEKPV_375_…_178_FK_179_…_236_NNI_238_. Notwithstanding the improbability of a match involving the tripartite anagram sSgII-9 + ‘sSgII-2’ + ‘sSgII-3’ being anything other than happenstance, it has been hypothesized that the EPL001 sequence is a true and meaningful reflection of sSgII structural reality, arising from the Edman machine reading *nonsequentially*

(Hart et al, 2022). In other words, EPL001 is to be found encrypted within sheep secretogranin II. The number of anagrams of a 14mer such as EPL001 is 14! = 87.18bn. Even though sSgII-14 is discontinuous (9+2+3), it is not a stray anagram of EPL001: it is an informatively constrained light shuffle. The sSgII-14 residues are a patterned interleaving of the EPL001 residues in regard to the front half of EPL001 (1-7, emphasized) and the back half (8-14): **ML**K**TG**E**KP**V·F**K**·NNI. The first doubleton, ML, has the initial M of EPL001. The next two shaded doubletons, TG and KP, are exact matches for doubletons within EPL001, albeit in reverse order: M**KP**L**TG**KVKEFNNI. (The probability that TG & KP occur in both sSgII-9 & the EPL001 equivalent 9mer is 1 in 1640, P = 0.0006, significant: Hart et al, 2022.) The last of the three shuffled-sequence modules, sSgII-3, comprises a key epitope component, NNI (Howlett et al, 2019).

sSgII’s second sorting domain is _367_**<u>MLKTGEKPV</u>**_375_. Motifs from this present in seven available ovine Edman sequences (S1 Table 1) are as follows: MLK (as ML/K, with MxLK in an eighth subsidiary sequence: see S1 Minor Sighting); TG (as TG or T/G, twice each, with MxxT in register in ‘Harwell 1’ of S1 Table 1); KP (thrice); and PV. In addition there is NNI (twice), which is present in sSgII outside its second sorting domain.

Suppose by way of a thought experiment that the sSgII-14 residues MLKTGEKPV·FK·NNI comprise a deck of 14 playing cards which when shuffled yields the EPL001 sequence MKPLTGKVKEFNNI. The 14×14 sequence grid of Fig. 10, plotting sSgII-14 along the top and EPL001 vertically, gives the residue matches, such that the shuffled ‘deck’ (EPL001) has the following face values: 1, 7, 8, 2, 4, 5, 11, 9, 3, 6, 10, 12, 13, 14. The EPL001 deck is said to have a ‘signature’ of four because it displays that many ‘rising sequences’ interleaved: <u>1 2 3</u>; <u>7 8 9 10</u>; <u>4 5 6</u>; <u>11 12 13 14</u>. An arithmetic ordering (i.e. 1-14) can be achieved simply by moving the 7-10 group between 6 & 11. The four groups of cards are patterned, 3-4-3-4. The starters, 1, 7, 4 & 11, are the sSgII-14 ‘peaks’ in the 14×14 grid, in order of decreasing height, with the other cards in each group being the associated ‘slopes’. To get to the strangely meaningful EPL001, imagine that the sSgII-14 deck is cut roughly into two halves which are placed on a table and corner-flicked together, effecting what is known as a riffle shuffle. In card playing the aim is to randomize the cards to defeat predictability. The sSgII-14 deck produces the EPL001 deck after (i) a rough halving (6/8), (ii) a riffle shuffle, (iii) a true halving (7/7) and (iv) another riffle shuffle. The transformation from sSgII-70 to EPL001 is as follows, with cuts represented by vertical lines and shuffles by arrows:

**Figure 10.**
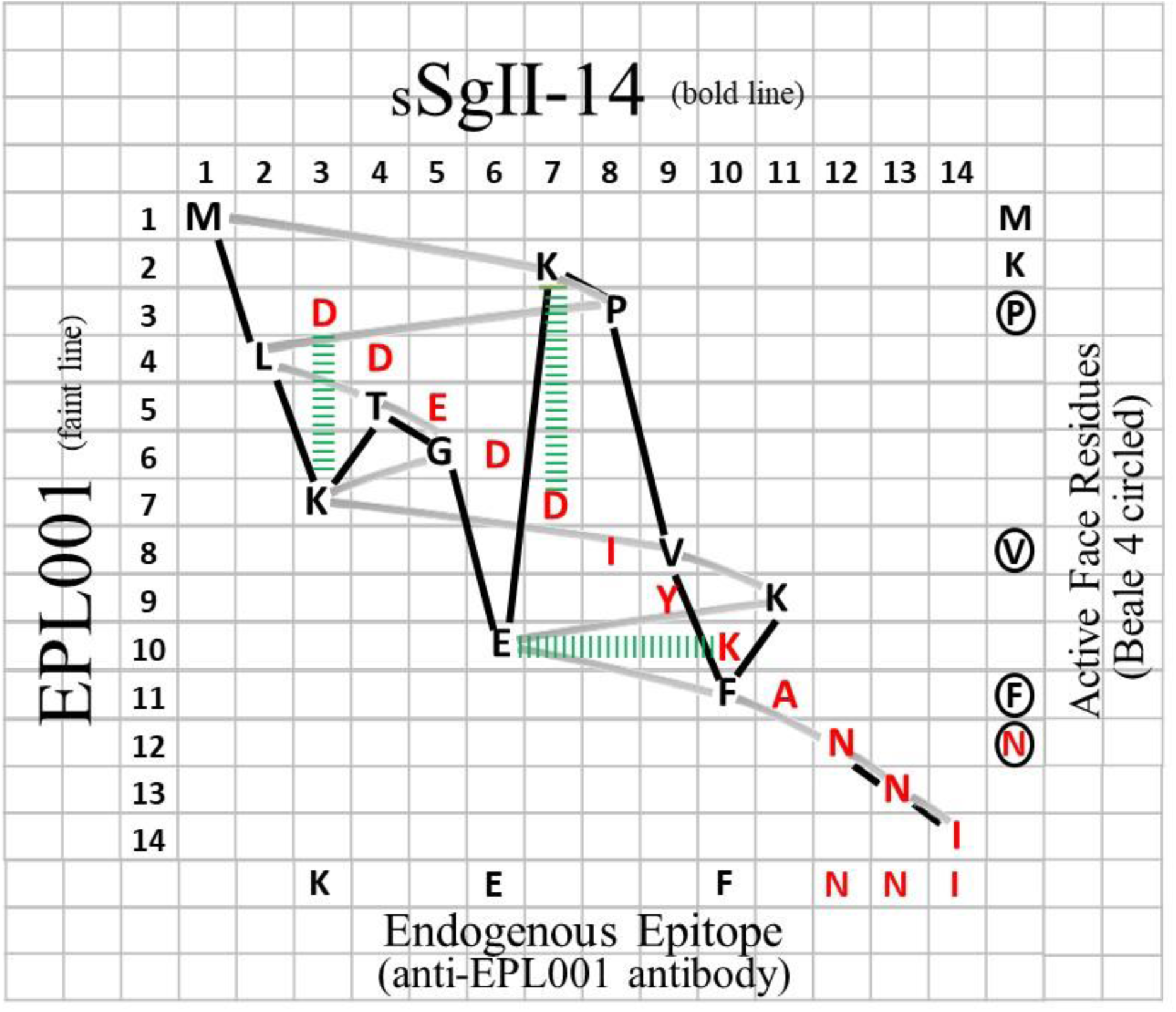
‘Magic grid’ modelling of sSgII-70 suggests overlapping ends. In black, N-terminal aa residues M1-K11 and in red C-terminal residues D59-I70. ‘sSgII-14’ (putative Edman N-terminal *input*) is plotted along the top against EPL001 (actual Edman N-terminal *output*) vertically. Following the bold line yields the aa sequence of sSgII-14 across the grid, column by column. The faint track reads out EPL001 downwards, row by row. For lysine grid placements see Hart et al, 2022. Green ladders represent potential salt bridges.

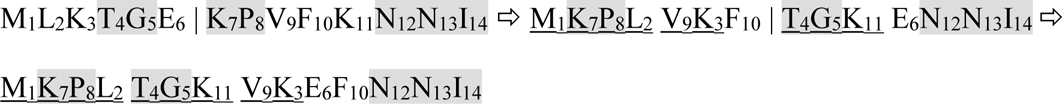

Adjacent shaded residues stay together throughout; adjacent underlined residues stay together through the second shuffle. *The adjacencies argue against a (far-fetched) chemical correlate to a riffle shuffling, with the economy of the transformation sSgII-14 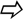 EPL001 supporting the view that the two sequences are non-randomly related.* With the signature of sSgII-14 by definition one, the signature of EPL001 after a series of shuffles could be from zero (14, 13, 12, 11, 10, 9, 8, 7, 6, 5, 4, 3, 2, 1) to seven, the latter in the form of doubletons: <u>13 14; 11 12; 9 10; 7 8; 5 6</u>; <u>3 4; 1 2</u>. A signature of four is what is typically obtained after two shuffles of a deck of 52 cards having equal independent placement probabilities, with eight shuffles and a plateau of 26 or so rising sequences representing adequate randomisation (Silverman, 2019). EPL001 thus has a high signature for two shuffles of a deck of 14, with its four rising sequences moreover covering all cards. What does all this mean? EPL001 is far from being a randomly ordered deck. The EPL001 Edman ‘card pick’ from sSgII-14 takes a very particular non-random form, connoting structural spirality in the manner of a spiral staircase.

IP/LC-MS, IHC, NNI-ome bioinformatics and sequence and shuffle analyses have vectored the project to sSgII and landed it on sSgII’s second sorting domain. If EPL001 is not a shuffled sequence from an Edman-challenging sSgII proteoform what is it? The N-terminus of a novel protein which by coincidence has granin-like characteristics in terms of IHC, acidity disclosed during anionex purification, heat resistance and stoichiometry? Or are the seven available EPL001-related Edman sequences in fact just gibberish?

For location purposes reference can be made to SgII in the rat. The second main sorting domain in rSgII comprises residues 366-380. It is situated in the wide terrain of primary structure between, on one side, the proteolytically derived peptides SN (184-216) & EM66 (219-284) and, on the other, a peptide product called manserin (529-568; Yajima et al, 2004). The EPL001/sSgII-14 anagram does not work in the rat or for that matter the human because of SgII-9 sequence variations: sheep MLKTGEKPV; rat MLK**A**GEKP**N**; human MLKTGEKP**N**. But given the *ovine* provenance of the EPL001 sequence these non-matches are immaterial.

The first of the seven available ovine Edman sequences is the ‘First Sighting’ (S1 Table 1). Derived from maximally purified (anionex) ovine ovarian follicular fluid this has the sequence MMxV?xPVG?GxFL. The First Sighting is a seeming prefigurement of EPL001 *and* ‘likely SgII relatedness’ *and* the identification of the protein Q9W2X8 as the fruit fly IHC antigen.

There is a rough correspondence in the form of M-V-F to EPL001: **<u>M</u>**KPLTGK**<u>V</u>**KE**<u>F</u>**NNI. The V in the First Sighting forms part of a PV doubleton, which is seen as such after M in the second sorting domain of sSgII (as the 9mer _367_**<u>M</u>**xxxxxx**<u>PV</u>**_375_) and in the homologue thereof in the fruit fly protein Q9W2X8 (as _1048_**<u>M</u>**xxxxxxxxxxx**<u>PV</u>**_1061_). (The **<u>P</u>**xxxx**<u>V</u>** of EPL001 on the other hand finds an in-register match with the ‘Beale 4’: xx**<u>P</u>**xxxx**<u>V</u>**/_L_xxF/_K_Nxx, with L and K as ‘under signals’: S1 Table 1.) The fruit fly’s homologue of SgII’s second sorting domain is within this sequence: _1038_PETLDFWQRE**<u>M</u>**ASKAG**<u>K</u>**TRNNQPVGS_1063_. Emphasized is the anchoring doubleton _1048_**<u>M</u>**xxxxx**<u>K</u>**_1054_, which matches _367_**<u>M</u>**xxxxx**<u>K</u>**_373_ in sSgII and _1_**<u>M</u>**xxxxx**<u>K</u>**_7_ in EPL001. The recognition of the importance of non-contiguous amino acid pairs that are evolutionarily connected has propelled understanding in the field of protein modelling (Crow, 2018). The M & K of sSgII’s **<u>M</u>**xxxxx**<u>K</u>** are the main peaks in the Fig. 10 grid comparing sSgII-14 and EPL001 and are analysed to be the first two residues to be read in Edman sequencing. The first and second sorting domains in the fly are self-similar in a way that is not the case in mammals. The match for the fly sorting domains is this: _45/1049_**<u>A</u>**x**<u>KAGK</u>**xxxx**<u>Q</u>**_55/1059_, with KAG a match for the second sorting domain of rSgII.

The 14×14 sequence grid of Fig. 10 compares sSgII-14 and EPL001, with sSgII-14’s 1-14 numbers along the top and EPL001’s 1-14 vertically. sSgII-14’s initial methionine is found at the start of EPL001, providing an M in the joint top left-hand corner of the grid. In the second column for sSgII-14 is leucine, M**<u>L</u>**KTGEKPVFKNNI, but this appears in the fourth row to match EPL001: MKP**<u>L</u>**TGKVKEFNNI. Matching the residues in this fashion provides for sSgII-14 an up-and-down zigzag and for EPL001 a side-to-side zigzag, representing the stepwise Edman reading order of the sSgII target molecule. A molecular model in silico of sSgII-9 has the same general shape as the 9mer part of the sSgII-14 line on the grid (S1 Fig. 3). EPL001 would appear to be an encoding with two aspects: of the complement of aa and of their relative positions in space. This explains why an antibody raised against EPL001 can see an sSgII proteoform.

### SgII-70

It has been asserted that there is a secreted 70mer polypeptide among the proteoforms of the sheep secretogranin II gene, *SCG2* (Hart et al, 2022). The deduced aa sequence of this 70mer polypeptide is represented in full in Fig. 11. This is sSgII-70, pronounced ‘sheep sig two seventy’. (The bovine equivalent is bSgII-70, rat rSgII-70, human hSgII-70.)

**Figure 11.**
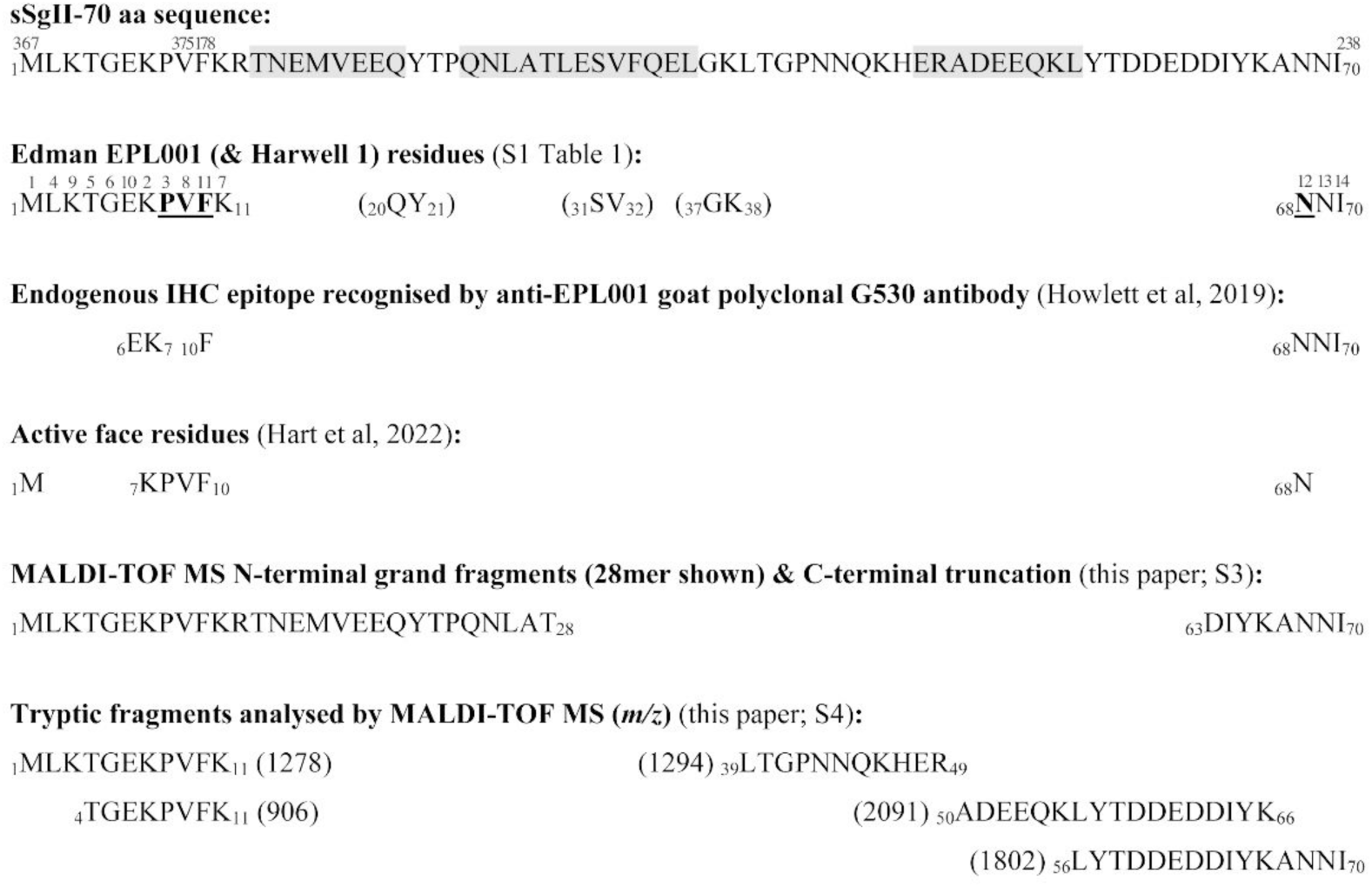
sSgII-70 primary structure deduction, with evidential support. sSgII-70 superscripts are positions in the sSgII master protein (Fig. 9). Helix predictions (shaded) by PSIPRED. EPL001 residues emphasized are anionex Beale 4. Edman superscripts denote spiralised reading order of residues within sSgII-70 to obtain EPL001. Bracketed Edman items (together with TG, unbracketed) appear thus in Harwell 1 (SEQ ID NO: 6 in S1 Table 1): MKV/I**<u>T/G QY S/V G?K?</u>**. A quadruple doubleton correspondence between Harwell I and sSgII-70 is unlikely to be due to chance (with Harwell 1 additionally sharing MK & TG with EPL001). Stepped-down gapped doubleton readings connote spirality.

Fig. 11 also displays for sSgII-70 the Edman-derived EPL001 residues and the order in which they were read (Hart et al, 2017); the IHC epitope residues recognised by the main anti-EPL001 antibody (Howlett et al, 2019); and the deduced active face of this inhibitory hormonal polypeptide (Hart et al, 2022). MS data from the present paper are also shown.

sSgII-70 would appear to be the product of peptide splicing (Howlett et al, 2019). This is the ‘9+61 contention’. Peptide splicing is distinct from RNA splicing. The sSgII prohormone can be modularised as ABCDE, as per Fig. 9. The 9 aa module is D, the 61 aa module B. If sSgII-70 is the product of *reverse splicing* from a single molecule of sSgII, then the processed product is DB. If sSgII-70 is the product of splicing from two molecules of sSgII, A1B1C1D1E1 + A2B2C2D2E2, then the possibilities are D1B2 & D2B1. (A chain arrangement of aggregated SgII molecules will be considered later.) Within the seven ovine Edman N-terminal sequences obtained (S1 Table 1) there is an M-F motif, seen once as _1_M-F_10_ and four times as _1_M-F_11._ This supports the 9+61 peptide splicing contention, with the 9-residue portion starting with M1 and the 61-residue portion with F10.

Further supporting the 9+61 peptide splicing contention for sSgII-70 is a re-evaluation of the evidence from tryptic digests, this time not driven by the fingerprint databases of already discovered and predicted proteins (see ‘Candidate 7500’). The analysis involves subjecting the proposed 9+61 arrangement of sSgII-70 to a tryptic digest in silico (Expasy PeptideMass) and searching for the predicted fragments in the MALDI data from the 13 trypsinisations attempted on Candidate 7500 material – and arguably finding examples, although de novo sequencing via MS/MS proved fruitless (S4). A tryptic digest spectrum is shown in Fig. 7 (with the peak list in S4 as Digest 5). The feedstock for this digest was an anionex fraction of ovine ovarian follicular fluid subjected to prior spin and gel filtrations. On the face of it this spectrum is discouragingly prolific, yet discernible in the understorey are four, maybe five, fragments of sSgII-70, as per the Fig. 7 legend. The peaks are in series in terms of intensity and a statistical analysis (of all five hits) supports a true correlation, with chi-squared P = 0.016, significant (S4 Table 3). Ovine MS candidate matches cover all trypsin-available residues in sSgII-70’s proposed primary structure (Fig. 11). Of 344 mass assignments in the ovine trypsinisation series a maximum of 25 (∼7%) might be sSgII-70 related (S4 Table 2). Crucial are two <u>integer</u> matches to sSgII-70’s proposed N terminus, at *m/z* **906**, _4_TGEKPVFK_11_, and **1278**, _1_MLKTGEKPVFK_11_. Both of these uphold the peptide spliced 9+61 junction at _9_VF_10_. The <u>integer</u> 1278 item was a singleton in the 11 ovine trypsinisations but the <u>integer</u> 906 signal appeared four times (= 36% prevalence), more than any item not a matrix cluster, trypsin autolysis product or keratin. The four digests producing 906 involved three different feedstocks purified in two different ways (S1 Babraham Method & Harwell Method) and analysed by two different MALDI mass spectrometers. Notably, 906 was present in ovine ovarian venous plasma but absent from jugular vein plasma from ovariectomised controls (S4 Digest 1), a result that is concordant with that for Candidate 7500, which is likewise OV+/OVX–. (The endogenous circulating antigen of the anti-EPL001 antibody is also OV+/OVX–, being present andabsent respectively in 3-30 kDa blood plasma ultrafiltrate of OV/OVX sheep). The MS for 906-producing Digest 1 was conducted by an external provider. The peak list (S4 Fig. 6) bears this comment: ‘…peaks consistent with autodigestion products of porcine trypsin, internal standards and commonly observed keratin contaminants or observed in the [gel] blank are noted’. The 906 signal was unannotated, as deemed analyte related.

The predicted mass of sSgII-70 is 8175.01 Da (Expasy Compute pI/Mw, using average isotopic masses). The protonated version of this, MH^+^, is 8176.02, with the addition of a hydrogen ion at 1.008 Da. Such a peak has not been observed with any modality of MS with any starting material, though in MALDI a **70mer** monomer-by-deduction has been seen in a putative quadruple homodimer (2+2+2+2) in a stored sample (see later, where is discussed S3 Fig. 10). In the Fig. 5 MALDI spectrum of EPL001 Gel Band 1 the main peak is at *m/z* 7583.64. Subtracting sSgII-70’s C-terminal –KANNI by peptide bond cleavage and *two* water molecules (with each H_2_O at 18.015 Da) provides a **65mer** <u>next-integer</u> match of 7582.36, with a difference of +1.28 and an error (difference as percentage of match) of 0.02% (Table 1). Remove a further water molecule and Fig. 5’s subsidiary peak at 7565.93 finds a **65mer** <u>next-integer</u> match in Table 1 at 7564.34, a difference of +1.59 and an error again of 0.02%. Alternatively, the same two peaks can be considered in terms of potential N-terminal losses from sSgII-70, again by peptide bond cleavage. For a match to the peak at 7583.64, subtracting MLKTG– and *three* waters provides a **65mer** match at 7590.28, difference –6.64 at an error of 0.09% (S3 Table 2). The peak at 7565.93 finds a **65mer** match *one* water below, with a difference of –6.34 at an error of 0.08% (S3 Table 2). What is true for the two peaks in Fig. 5 is true for all the 7-8000 peaks in ovine Figs. 2-5, taken together (n = 12, Table 1): MALDI data modelling based on C-terminal aa residue losses provides a snugger fit for the ovine data than N-terminal residue losses (S3). Seven of the 12 ovine peaks find <u>integer</u> or <u>next-integer</u> matches (58%) in the C-terminal truncation analysis, three (25%) in the N-terminal truncation series.

The superior data fit for *ovine* SgII-70 of C-terminal truncation over N-terminal truncation also holds true for the *bovine* series. Peaks of *m/z* 7-8000 (n = 13) are analysed from Fig. 4 (top panel, 3 peaks), Fig. 6 (8 peaks) & S3 Fig. 1 (2 peaks). There are three residue differences between the ovine and bovine SgII-70 aa sequence predictions: M16 in ovine Fig. 11 becomes bovine I16; N44 becomes S44; and H47 becomes R47. The predicted *m/z* scores are thus different throughout for the 7-8000 peaks. *C-terminal truncation* in the bovine analysis (S3 Table 3) delivers a mean deviation of **2.99** with a mean error at **0.04%**; two <u>integer</u> or <u>next-integer</u> matches (**15**%); ∑Observed (n = 13) 97097/∑Matches 97117 x100 = **99.98**%; and a chi-squared P = 0.027, significant. *N-terminal truncation* relating to the same bovine peaks (S3 Table 4) delivers in contrast a mean deviation of **5.13** with a mean error at **0.07%**; one <u>next-integer</u> match (**8**%); ∑Observed (n = 13) 97097/∑Matches 97073 x100 = **100.03**%; and a chi-squared P = 0.057, ns.

The MS analysis amounts to a C-terminal sequencing that upholds the concluding part of SgII-70 for both ovines and bovines as _63_DIYKANNI_70_. (The same conclusion holds for the pig, on the basis of a sole available spectrum, S3 Fig. 2, in which an apparent pSgII-70 **68mer** peak at *m/z* 7875.58 finds a predictive <u>next-integer</u> match at 7876.72, with *four* water losses, and an apparent **61mer** at 7088.26 finds a match at 7092.89, with *two* water losses.) Consider the upper panel of Fig. 4. The difference between the two peaks, both relating to SgII-70 in *bovine* ovarian follicular fluid, is equivalent to an alanine residue. The 7643.93 item relates to a **67mer** ending in –DIYKA; the 7573.19 item to a **66mer** ending in –DIYK (S3 Table 2). During one purification campaign using *ovine* jugular vein plasma as feedstock in a Babraham style approach (S1), with 1D SDS-PAGE followed by whole gel elution, two prominent peaks were recorded using MALDI-TOF MS (Ciphergen, University of Leeds, UK; personal communication, Julian Hiscox) at *m/z* 7191 and 7565. There are <u>next-integer</u> matches to these in Table 1 at 7190 (**62mer** with *two* water losses) and 7564 (**65mer** with *three* water losses).

A data fitting exercise based on C-terminal truncation and water losses works in tying MALDI observations to ‘predictions’ (meaning *matching a model*, i.e. fitting into an analytical framework, given a certain *m/z* observation, not *forecasting*), but is disconcerting. Molecular losses from a small protein would normally require exposure to extreme chemical or physical events or enzymatic attack, none of which is relevant in the present case. How does the heterogeneity of Candidate 7500 in MALDI reflect the situation indicated by other approaches? The aa motif NNI is the C-terminus of both EPL001 and seemingly sSgII-70. For EPL001 to be obtained by the Edman machine sSgII-70 must be present in its entirety, yet no stand-alone MS peak has been seen corresponding to an intact **70mer**, as pointed out earlier. Now consider Fig. 5 once more. This mass spectrum relates to Gel Band 1 which in a prior purification yielded the EPL001 Edman sequence MKPLTGKVKEFNNI. The three peaks in the mass spectrum can be analysed as follows in terms of sSgII-70 in Table 1: 7930.24 (**69mer** lacking –I) and 7583.64 & 7565.93 (**65mers** lacking –KANNI). It is not possible to derive the 14-residue EPL001 sequence from any of these truncated forms, in which the C-terminal NNI is diminished or lacking altogether, so it seems that C-terminal truncation is an artefactual aspect of the MALDI campaign. Sheep serum in bulk subjected to anionex and western blotting with an anti-EPL001 antibody produced mono-bands at 0.2M and 0.8M NaCl at ∼7 kDa (S1 Fig. 4). For this to be the case the C-terminal partial epitope –NNI must be present. In other words, some amount of SgII-70 must be present in its entirety, in line with the Edman EPL001 data. The anionex purification process used was the same as in the MS investigations, but MALDI-style heterogeneity was not seen. Laddering *was* seen however in a western of aqueous extract of rat hypothalamus (Hart et al, 2017). The ‘rungs’ on the ladder are too close for whole-factor multimerization to be at issue and anyway for the rungs to be visualised by the anti-EPL001 antibody the C-terminal NNI must be present. So, MALDI-style C-terminal truncation is not relevant to the observed laddering, which has been attributed to splice variants in the rat not seen in the sheep. An aqueous extract of ovine jugular vein 3-30 kDa upstream precipitate was analysed by SDS-PAGE (text after S1 Fig. 10). Several Coomassie-stained bands above 6 kDa showed the presence in MALDI of Candidate 7500, with no or a weak correlation of MW with placement up the gel. It is concluded that MS heterogeneity in the range *m/z* 7-8000 relating to C-terminal truncation of Candidate 7500 is exclusively associated with the MALDI campaign.

What is to be made of the annotated peaks below *m/z* 7-8000 in the MALDI spectra of Figs. 2-4, there being none such in Figs. 5 & 6? These Ovine Minor Peaks can be submitted to the same sSgII-70 heterogeneity analysis as peaks in the 7000s, involving losses of C-terminal residues and water losses. This uses S3 Table 1, a version of Table 1 extended downwards through the mers. The results are as follows: (i) Observed *m/z* 6109.35 vs Match 6102.85 (**54mer** with *eight* water losses); (ii) 6070.58 vs 6064.79 (**53mer** with *three* water losses); (iii) 5655.12 vs 5655.44 (**50mer** <u>integer</u> match with *five* water losses); (iv) 5070.90 vs 5069.78 (**45mer** <u>next-integer</u> match with *three* water losses); (v) 4785.16 vs 4791.51 (**43mer** with *three* water losses); (vi) 4755.85 vs 4755.48 (**43mer** <u>integer</u> match with *seven* water losses); & (vii) 4748.66 vs 4749.47 (**43mer** <u>next-integer</u> match with *seven* water losses). In summary: ∑Observed (n = 7)/∑Matches x100 = 37195.62/37189.32 x100 = **100.02%**; chi-squared P = 0.02, significant (S3 Table 9).

Of the seven Ovine Minor Peaks just analysed four occurred in the mass spectrum in the lower panel of Fig. 4. The material there was a solid ‘upstream precipitate’ that formed on the 3 kDa membrane during a bulk 3-30 kDa ultrafiltration of ovine jugular vein EDTA plasma (S1 Sheffield Method). Bacterial contamination was obvious in terms of discolouration and confirmed by Edman sequencing. Can it be true that the only thing that flew in MS from this untreated sludge was the sought-for sSgII-70? Apparently, yes. The use of sodium azide as an antimicrobial agent reduced but did not eliminate the upstream precipitate, which now appeared white. Separation of this uncontaminated precipitate by SDS-PAGE saw EPL001 Gel Band 1 being obtained by electroelution to provide the uncluttered mass spectrum of Fig. 5. An alternative approach was to use solvent extraction of gel bands to provide upstream precipitate for MALDI (S3 Fig. 5). As in Fig. 5, the base peak (i.e. the most prominent peak, taken as 100%, in an intensity plot of relative abundance) in S3 Fig. 5 at 7579 is analysed via Table 1 to be a **65mer**, i.e. sSgII-70 lacking –KANNI, with *two* water losses. But whereas Fig. 5 is uncluttered, the S3 Fig. 5 spectrum resembles the lower panel of Fig. 4, except that the minor peaks are higher in relation to the base peak and there is a raggedness to the spectrum that may be due to the presence of SDS, absent from the Fig. 4 preparation. There are multi-hundred stepped peaks which can be analysed by reference to S3 Table 1 as N-terminal Grand Fragments, just like the Ovine Minor Peaks. A sub-8000 analysis is as follows: 3791 (**33mer** match 3789), 4607 (**40mer** match 4613), 5998 (**53mer** match 5992), 6741 (**59mer** <u>integer</u> match 6741), 7579 base peak (**65mer** match 7582) & 7924 (**69mer** match 7919): ∑Observed (n = 6) 36640/∑Matches 36636 x100 = **100.01%**.

Apparent N-terminal Grand Fragmentation of a uniform kind was seen when anionex HPLC fractions of ovine jugular vein EDTA blood plasma were analysed by MALDI-TOF MS (M-Scan, Wokingham, UK), after pooled samples were desalted using Sep-Pak C18 cartridges (S3 Fig. 8 legend). Apart from discounted Candidate VI, fibrinopeptide A, in the form of peaks in the *m/z* 1400s subject to MS/MS sequencing, the main finding was an array of peaks in the *m/z* 3000s. As an example, one MALDI spectrum (S3 Fig. 8) is bare of significant peaks save for a cluster in the 3000s, topped by signals at *m/z* 3196, 3179 & 3161. In S3 Table 1, the downwardly extended version of Table 1, these ions find matches on the sSgII-70 **28mer** tier at 3195 (*four* water losses), 3177 (*five* water losses) & 3159 (*six* water losses). When the matrix was CHCA and the instrument was in reflectron mode what were mainly seen were sSgII-70 **28mers**. With sinapinic acid as matrix and the MS instrument in linear mode what were mainly seen were sSgII-70 **29mers**. (By way of comparison, an anionex sample of ovine jugular vein EDTA plasma at Babraham provided major peaks in MALDI at 2420, being a putative **21mer** having a match in S3 Table 1 at 2415, and 2342, being a putative **20mer** with an <u>integer</u> match at 2342, the former obtained with CHCA as matrix, the latter with sinapinic acid.) The full analysis of the M-Scan data takes in 23 peaks in the *m/z* 3000s seen in 10 of 16 mass spectra (S3 Table 10) and involves 13 <u>integer</u> and <u>next-integer</u> hits (57%) to S3 Table 1 values such that ∑Observed (n = 23) 74406/∑Matches 74402 x100 = **100.01%**. A Babraham spectrum of late-eluting anionex Candidate 7500 from ovarian follicular fluid has peaks supportive of the contention that ovine MALDI peaks in the *m/z* 3000s are sSgII-70 N-terminal Grand Fragments: 3233 (**28mer** S3 Table 1 match 3231), 3553 (**32mer** <u>next-integer</u> match 3552) & 3924 (**34mer** match 3917) (S3 Fig. 11). Solvent extracted gel bands of upstream precipitate yielded half-sized peaks, as follows: main peak at 7572 (**65mer**), with several peaks including one at 3787 (7572 ÷ 2 = 3786, **33mer** match 3789); and main peak at 7830 (**68mer**), with a sole other peak at 3916 (**34mer** <u>next-integer</u> match 3917, with 7830 ÷ 2 = 3915,) (see text after S1 Fig. 10, ‘Extracted bands of contaminated material’). Purification Candidate IV was ‘Luteal phase entity’ at *m/z* 3450 (S1). It was inactive in vitro and in vivo. It is analysed to be an N-terminal Grand Fragment of sSgII-70: **30mer** match 3453.

The half-sized ‘Sheep 3000’ N-terminal Grand Fragments result has been replicated in the cow. A late-eluting anionex fraction of *bovine* follicular fluid was subject to MALDI analysis after five years storage, with the instrument in linear mode using sinapinic acid as the matrix (S3 Fig. 9, interpreted using S3 Table 3). The only significant peaks were a cluster in the *m/z* 3000s mainly representing putative bSgII-70 **29mer** Grand Fragments, as per the use of sinapinic acid in the sheep. ∑Observed (n = 5) 16339/∑Matches 16337 x100 = **100.01%**, as in the sheep.

Dimerization is discernible, where ‘dimer’ refers to factor fragments doubled up to form *homodimers* (αmer + αmer) and *heterodimers* (αmer + ß mer), with the ‘mer’ of di*mer* meaning ‘peptide’ and the ‘mer’ of α*mer* and ß*mer* denoting residues in α or ß numbers. (Dimer is NOT a reference to a **2mer**, which would be an isolated factor fragment having two aa residues, a ‘two-mer’.) In Fig. 4 the peak at 9509 is a ‘Superheavyweight’, defined as an ion above the predicted MH^+^ mass of sSgII-70 at *m/z* 8176. This ion at 9509 is twice the mass of the lowest peak, at 4755 (**43mer** S3 Table 1 <u>integer</u> match 4755, with *seven* water losses). N-terminal Grand Fragment *homodimerization* (αα) is implied: MH^+^ at 4755 + M at 4754 = 9509, <u>integer</u> match to Observed. A Superheavyweight in S3 Fig. 5 at 8383 is roughly the sum of the two lowest peaks, this time implying N-terminal Grand Fragment *heterodimerization* (αß): 3791 (**33mer** match at 3789) + 4607 (**40mer** match at 4613) = 8398. S3 Fig. 5 is the MALDI spectrum for solvent extracted EPL001 Gel Band1 (see text after S1 Fig. 10, ‘Uncontaminated gel’). Carrying on up the MW gel ladder, the next two bands boast items at 8935/8933, which are twice the size of items in the same samples at 4466/4468 (**40mer** <u>next-integer</u> match at 4469), potentially indicating once more N-terminal Grand Fragment homodimerization producing a Superheavyweight: MH^+^ match at 4469 + M at 4468 = 8937. A conclusion is that the peaks in the lower panel of Fig. 4 and in the S3 Fig. 5 spectrum are exclusively sSgII-70 related and rampantly artefactual, including in the matter of N-terminal Grand Fragment dimerizations producing Superheavyweights. The mers in Fig. 4’s lower panel are as follows, with S3 Fig. 5 mers in brackets: - (**33**), **43** (**42**), **50/53/54** (**53**), **62** (**59**), **65/66** (**65**) & **67** (**69**). These steppings hint at patterned grand fragmentation.

The bioactivity of purified samples can be lost even after a few days of storage, indicating compromised molecularity (see shortly, slopes analysis, **7mer**). Anionex fractions of ovine plasma have been subject to analysis after five year’s storage (S3 Fig. 10). Fraction 24 (of 30) yields a picture via MALDI of sSgII-70 **19-64mers** in the form of N-terminal Grand Fragments up to items in the 7000s, together with potential homodimers (x2) and multi-homodimers (2+2, 2+2+2, 2+2+2+2). The base peak is 14615, a putative homodimer: **63mer** MH^+^ monomer match 7306 + M of 7305 = 14611. Then there are peaks stepped up in mass but stepped down in intensity at 28831 (**62mer** MH^+^ monomer <u>integer</u> match 7208 + M of 7207 x3 = 28829), 43227 (**62mer** MH^+^ monomer match 7208 + M of 7207 x5 = 43243) & 64974 (**70mer** MH^+^ monomer <u>integer</u> match 8121 + M of 8120 x7 = 64961). The analysis of this spectrum thus involves five sub-8000 non-dimeric peaks (Observed vs Matches) plus ten Superheavyweight putative homodimers (Observed) *or* ten Superheavyweight-homodimer-matches-by-calculation (Matches), thus: ∑Observed (n = 15) 310596/∑Matches 310533 x100 = **100.02%**.

Slopes on the left-hand sides of the Figs. 2 & 3 spectra and many others in S3, often providing base peaks, call into question MALDI instrument set-up and operation. The items in these slopes are not accounted for by matrix ions, which tend to be in the *m/z* 800s (Keller et al, 2008), with the low mass gate mostly set anyway at 1000 Da. Slopes are seen in MALDI intensity plots of relative abundance involving Candidate 7500, though not always (e.g. Fig. 4), but slopes are *not* seen in counts plots, though the same peaks are present (S3 Fig. 14a & b). An intensity plot of an ovine plasma anionex fraction (S1 Sheffield Method) is presented in a supplementary file (S3 Fig. 13d). Its most prevalent two peaks, at the top of the ramp, can be analysed in terms of sSgII-70: 1132 base peak (**10mer** S3 Table 1 <u>next-integer</u> match 1133), 1059 (**10mer** match 1061). Peaks at 1133 and 1059 are similarly available from a separate ovine plasma anionex fractionation (S1 Harwell Method). With a counts plot and a low mass gate set at 600 Da there are no other annotated peaks and the baseline is horizontal (S3 Fig. 14a). An attempt to bulk up analyte involved combining and concentrating eight anionex fractions of ovine blood serum containing Candidate 7500 (S1 Babraham Method). No MALDI peaks in the *m/z* 7000s were evident two days later. Three peaks in the 2000s were seen instead, with a ramp on the left-hand side (S3 Fig. 12). The 2000s are interpreted as sSgII-70 Grand Fragments: ∑Observed (n = 3) 7430/∑Matches 7434 x100 = 7430/7434 x100 = **99.95%**. The analysis of the highest intensity peak is as follows: 699 base peak (sSgII-70 **7mer** <u>integer</u> match 699, with *five* water losses). Analysing the spectra alluded to above in terms of sSgII-70: Fig. 2, 1027 (**10mer** match 1025) & 1149 base peak (**11mer** match 1153); Fig. 3, 1130 (**10mer** match 1133) & 1147 base peak (**11mer** match 1153). An aggregated analysis can be offered on the basis of dually decremented sSgII-70 of ramp peaks in six ovine spectra, all intensity plots (S3 Fig. 8, S3 Fig. 11 & S3 Fig. 13a-d): ∑Observed (n = 35) 87503/∑Matches 87442 x100 = **100.07%**. Concluding, the slopes on the left-hand side of *ovine* MALDI intensity plots are Candidate 7500 fragment ramps.

An aggregated analysis of *bovine* ramp peaks can be offered on the basis of dually decremented bSgII-70 in a pair of spectra, both intensity plots (S3 Fig. 1 & S3 Fig. 9): ∑Observed (n = 11) 14960/∑Matches 14948 x100 = **100.08%**.

At each step across Table 1 there is the loss of a mass equivalent to neutral H_2_O from an unhydrated (water-free) sSgII-70 C-terminally decremented polypeptide. Water gaps are annotated in the mass spectrum of Fig. 6 relating to bSgII-70. Ovine systemic plasma was subjected to spin and gel filtration, followed by anionex (S1 Babraham Method). Fraction 20 showed a single peak in MALDI at *m/z* 7495 (S3 Fig. 6), betokening an sSgII-70 **65mer** with *seven* water losses, according to Table 1: match 7492. Reread within minutes using the same matrix, this fraction again yielded only a single peak, this time at 7550 (S3 Fig. 7), still corresponding to a **65mer** but now with *four* water losses (match 7546). Why the difference? The 7495 item was detected when the instrument was in continuous mode (i.e. with immediate lasering), the 7550 item when it was in delayed ionisation mode. A *reduction* in water losses due to a minor deferral of lasering is machine artefactuality of an intriguing kind.

The dually decremented MS data model for SgII-70, featuring C-terminal aa truncation of the **70mer** together with water losses, surprises by working for almost-intact monomers (**60mers**, Table 1) down to minor fragment base peaks, all N-terminal, as well as for Superheavyweight heterodimers (αß), homodimers (αα) and multi-homodimers (αα x n), likewise with all dimers involving dually decremented N-terminal monomer fragments. In the IP/LC-MS antigen capture campaign that took the project to ‘Likely SgII related’, protean inscrutability was trammelled by the use of feedstock subject to formalin fixation with antigen retrieval.

### Taking stock

Hypothesis H1 is that an inhibitory gonadal influence on tissue masses IS accounted for by the factors described (Candidate 7500, EPL001 & SgII-70). Hypothesis H2 is that an inhibitory gonadal influence on tissue masses IS NOT accounted for by the factors described (Candidate 7500, EPL001 & SgII-70).

The chain of logic behind this paper – from Candidate 7500 through the (machine-artefactual) EPL001 sequence to sSgII-70 and back to Candidate 7500 – is shown in Fig. 8 and available in tabular form as S6 Table 1. Although peculiar in finding myriad MS peaks of Candidate 7500 to be related to a single bioactive polypeptide – *E pluribus unum –* and the canonical EPL001 N-terminal sequence MKPLTGKVKEFNNI to be lightly mixed up within the neuroendocrine prohormone sSgII, mostly in its second sorting domain, as MLKTGEKPV·FK·NNI (Fig. 9), these interpretations are upheld statistically and support H1 by delivering potential insights (S6 Table 2). Chief among these is the deduction of sSgII-70’s active hormonal face as **<u>M·KPVF·N</u>** (Fig. 11), with that of human and rat being **<u>M·KPNF·N</u>**. These realizations have permitted the synthesis of hexapeptide mimetics displaying relevant bioactivities in the form of antiorganotrophism (based on fewer smaller cells) and reproductive modulation (Hart et al, 2022).

Candidate 7500 is present in blood plasma from ovary-intact sheep and absent in that from OVX ewes, with the former material bioactive in vitro, the latter not. ‘OV+/OVX–’ is true too of the circulating antigen of an anti-EPL001 antibody and also of an MS tryptic fragment at *m/z* 906 (_4_TGEKPVFK_11_) predicted from the trypsinisation in silico of sSgII-70. If the 906 item were instead OV+/OVX+ then H1 would be subject to a Baconian exclusion of hypothesis (Platt, 1964). Presence of 906 in material from OVX ewes would be incompatible with its being related to a secreted hormone which circulates from a gonadal source, while having, on the basis of IHC, a conjectured paracrine hypothalamic manifestation too. (Note that oestradiol circulates from the gonads but is also made in the hypothalamus.)

No findings unequivocally discount H1, no findings unequivocally support H2. H2 explains nothing, leads nowhere and is rejected. H1 is supported by different lines of evidence – with better than triangulated support for ‘OV+/OVX–’ (Fig. 8) and triangulated support for ‘EPL001 does not exist in nature’ and ‘Likely SgII related’ – co-identifying Candidate 7500, EPL001 & SgII-70 with each other and with the sought-for inhibitory gonadal hormonal influence on tissue masses, micrin.

### Summation

The White Queen in *Through the Looking-Glass* claims to be able to believe six impossible things before breakfast. The present article asks less of the reader, providing (a) MALDI data fitted within a unitary sheep and bovine ‘SgII-70’ model based on peptide splicing from secretogranin II and featuring extravagant artefactual molecular losses and dimerizations (factor fragments doubling up) of a kind not usually associated with MALDI investigations; (b) wayward chemical sequencing in a manner not reported before (‘Edman Nonsequentialism’), apparently due to SgII-70 undergoing a spiralised depolymerisation in the Edman reaction chamber, notably providing a baffling 14 aa N-terminal sequence denoted ‘EPL001’; and (c) a polypeptide aa sequence decryption from SgII using bespoke bioinformatics and bioactivity profiling, which discloses the hormonal active face of its derivative SgII-70 and permits the production of synthetic hexapeptide mimetics thereof; with, along the way, (d) an ad hoc tryptic digest analysis upholding the SgII-70 identification, in particular via an ion at *m/z* 906 which is an <u>integer</u> match to a predicted tryptic fragment of sSgII-70, and (e) a rarefied epitope mapping exercise based on IHC, identifying the binding site endogenously of an anti-EPL001 antibody as SgII-related. That’s five.

The current study – with its evidential triangulations and better, replicated results in separate laboratories by different researchers and multi-modal statistical significance – has been an exercise in classical endocrinology, with added cryptanalysis. Genetic approaches involving expression patterns, mutations, knock-outs and so on have to an extent superseded the isolation of factors by biochemical enrichment, given that fruit fly morphogens for example are present in trace amounts and assays to detect them in biological extracts tend to lack reliability (Nüsslein-Volhard, 2004). Yet direct analysis of the secretome is still imperative, to characterise what’s actually being made and exported. SgII-70 has been panned from the downstream mysterium.

In the hunt for a postulated tissue-mass inhibiting gonadal hormone there emerges Candidate 7500 from physicochemical purifications of ovine and bovine blood and ovarian follicular fluids. Candidate 7500, ostensibly a polypeptide, displays an enigmatic *m/z* 7-8000 heterogeneity in MALDI, while being undetectable in other MS modalities. Associated cell-killing activity in vitro is protease and heat resistant. With difficulty aa sequence data are extracted. The 14-residue Edman N-terminal sequence EPL001 is glimpsed in maximally purified ovine plasma anionex fractions as the Beale 4, xx**<u>P</u>**xxxx**<u>V</u>**xx**<u>FN</u>**xx, before being apprehended more fully in upstream material as MK**<u>P</u>**LTGK**<u>V</u>**KE**<u>FN</u>**NI (S1 Table 1) –apprehended by bioinformatics and molecular biology as inscrutable. Inscrutability is unscrewed when from the use of an anti-EPL001 antibody in IP/LC-MS there emerges the concept of ‘Likely SgII related’, a reference to the intracellular neuroendocrine prohormone secretogranin II. This result is triangulated with the addition of evidence from IHC and bespoke bioinformatics. Micrin is SgII related, it is surmised, in a ‘Let there be light!’ moment. EPL001 is decrypted as a mixture of both ends of a 70 aa polypeptide derivative of SgII, sSgII-70. The 14×14 plot of Fig. 10, showing Edman *output* (EPL001) vertically versus putative sSgII-70 *input* across the top, is deemed a ‘magic grid’, by dint of conjuring up apparent structural insights. The deduced sequence of SgII-70 is dually decremented in an MS datafitting model involving C-terminal truncation (i.e. loss of C-terminal aa residues as a result of peptide bond breakage) and water losses, upholding the C-terminal sequence of sSgII-70 & bSgII-70. The heterogeneity of Candidate 7500 in MALDI is deduced to include the production from sSgII-70 of N-terminal Grand Fragments down to a **7mer** and even Superheavyweight ions in the form of fragment heterodimers (αß), homodimers (αα) and multi-homodimers (αα x n). Breathless stuff.

Circular explanatory scheme, circular reasoning? No. Candidate 7500 anionex fractions, EPL001 in the form of a 14mer synthetic peptide, as well as hexapeptide mimetics of SgII-70’s deduced active face: all three have independently delivered inhibitory bioactivity readouts (Fig. 8). This is yet another triangulated result.

What is to be made of the observed heterogeneity in MS? MALDI is a soft ionisation technique, with laser energy imparted to target molecules via an organic matrix. It is an analytical method based on intact mass. A contrast is tandem mass spectrometry (MS/MS), which technique does indeed feature peptide bond breakages, molecular losses and the generation of fragment ions. In MS/MS peptide ions are separated in a mass analyser according to *m/z* then they are further fragmented and ionized during a collision-induced dissociation step. Comparable results in MALDI would involve post-source decay – of a kind never before reported with this technique and hence not favoured as an explanation here.

Seen in MALDI spectra, according to the analysis, are N-terminal **b**-ions (to use MS/MS fragmentation terminology) of SgII-70, as well as dimers thereof, all subject to the dual decrementation of C-terminal truncation and neutral water losses. With a lack of chemical, physical or enzymatic assaults on the target polypeptide, molecular decrements are perplexing, though it is possibly relevant that acidic conditions are a feature of both MALDI plate set-up and Edman degradation. In MS/MS water losses from peptides can involve the loss of carboxylic acid oxygen, side-chain hydroxyl oxygen or peptide backbone oxygen (Ballard & Gaskell, 1993), occurring most prevalently from the hydroxylated side chains of S & T, with such losses also associated with G, D, Q & N (Sun et al, 2008). Another explanation for the water decrements overall can be discounted as implausible, as encompassing paucity and plethora. This would be based on transester condensation crosslinking of a variable kind (0-8), involving the hydroxyl groups on the side chains of S, T & Y and the carboxyl groups of the side chains of D & E, releasing water. The view taken here is that SgII-70 loses C-terminal parts of itself plus water during processing and storage and due to MALDI ‘plate tectonics’ (choice of matrix and immediate or delayed lasering), rather than as a result of MS post-source decay. It can be claimed that SgII-70 has been purified to homogeneity on the basis that in anionex fractions the only other entity identified (by *m/z* ratio and Edman degradation) was fibrinopeptide A, Candidate VI, which was discounted as contra-active and which was seen only in a tiny minority of spectra (e.g. S3 Fig. 14b).

Three manifestations of sSgII-70 can be discerned in the MALDI data: (i) Candidate 7500 (= peaks in the m/z 7-8000 range showing dual decrementation to provide **60-70mers**); (ii) N-terminal Grand Fragments (= fragments showing major losses from the C terminus, plus water losses), notably producing a stepped grand fragmentation pattern of **20mers**, **30mers**, **40mers**, **50mers** and **60mers**, as well as smaller base peaks; and (iii) Dimerization (= the provision of αα fragment homodimers and αß fragment heterodimers by the doubling up of dually decremented monomer fragments, such dimerization not being seen in electrophoresis).

Referring to (i) and (ii), bulked-up anionex fractions of Candidate 7500 provided a MALDI spectrum after a couple of days storage devoid of Candidate 7500 but with **19-21mer** peaks in the 2000s and a base peak at *m/z* 699 that was an <u>integer</u> match to a predicted **7mer** N-terminal fragment of sSgII-70 (S3 Fig. 12). Yet this latter match to the sSgII-70 MS data model involves a credulity-stretching *five* water losses. This, from a **7mer** of deduced sequence MLKTGEK. Assuming the 699 result is not a freak coincidence, it is hard to understand how the water losses part of the MS model can go on working when there is only one-tenth of the aa complement present, unless all the water losses in Table 1 and its extended form, S3 Table 1, come from the N terminus. Other grand fragmentations have been described in regard to the use of reverse phase chromatography after anionex and separately relating to consecutive MALDI evaluations of an anionex fraction in different laboratories (‘Candidate 7500’).

Referring now to (iii), Dimerization, this as an artefact in MS relates to ionisation of target molecules, generating (non-covalent) electrostatic bonds between monomers, with shared charge. Fig. 4 sports both a dually decremented sSgII-70 fragment monomer (a **43mer** <u>integer</u> match at 4755, with *seven* water losses) and its homodimer (at an <u>integer</u> matched 9509 to Observation), doubly dual decremented. So this is the simultaneous presence in the same spectrum of a decremented Grand Fragment and a homodimer thereof. A five-year stored sample of ovine anionex plasma ostensibly delivered fragment homodimers in the pattern 2, 2+2, 2+2+2, 2+2+2+2 (S3 Fig. 10). The component monomers are analysed to be dually decremented. Leaving aside the molecular alterations and the locus in the analytical process where these might occur, there is factor identification. There are <u>integer</u> and <u>next-integer</u> matches to the prediction of an SgII-70 dual decremented MS data model across the factor-fragment size range, **7-70mer**, with correspondences to the model for data groups, including dimers, in all cases at around 100%, amid the achievement of high degrees of chi-squared significance as between Observed and Matches. The model works. MALDI is deemed to have read true throughout, detecting SgII-70 faithfully, though in a manner requiring careful analysis.

Regarded as a pharmacon (bioactive molecule) SgII-70 has as a conserved pharmacophore (active face) **<u>M·KPV/NF·N</u>**. This has been deduced from studies in vivo and in vitro using synthetic peptides (Hart et al, 2022). The sSgII-70 active site **<u>M·KPVF·N</u>** is found in ovine derived EPL001 as follows: **<u>MKP</u>**LTGK**<u>V</u>**KE**<u>FN</u>**NI. This realisation accounts for (i) the relatively weak inhibitory properties of EPL001; (ii) the ineffectiveness in preventing EPL001’s inhibitory activity of alanine substitution by sector (N terminus, Centre Section, C terminus); (iii) the inhibitory activity of both N-terminal and C-terminal EPL001-related hexapeptides; and (iv) the anomalous inhibitory activity in vivo and in vitro of a scrambled-sequence control peptide (EPL030), which happened to have in its sequence **<u>PVF</u>**, three of the Beale 4 (S1 Table 1), and an appropriately spaced **<u>K</u>-<u>P</u>** doubleton: KL**<u>K</u>**MNGKNIE**<u>PVF</u>**T. In terms of drug discovery there has been a journey from the 14mer parent compound EPL001 to lead candidate hexamer MKPVFN (EPL601). EPL001, representing only one-fifth of sSgII-70’s complement of aa, is receptor active by virtue of its being an amalgam of the two ends of sSgII-70, with 56 intermediate residues (80% of the total) absent.

EPL001 cannot be found using tryptic digestion, molecular biology and bioinformatics because a source protein with this N-terminal aa sequence doesn’t exist in nature. This is a triangulated negative. How in principle can an antibody raised against a synthetic linear sequence of 14 aa see a different primary sequence endogenously, with the two sequences being anagrams of one another? So, that is an antibody to _1_MKPLTGKVKEFNNI_14_ (EPL001) binding to _1_MLKTGEKPVFK_11_**·**_68_NNI_70_ (sSgII-70), with epitopes in the form of _9_**<u>KEFNNI</u>**_14_ and _7_**<u>KE</u>**_6_**·**_10_**<u>F</u>·**_68_**<u>NNI</u>**_70_, respectively (Howlett et al, 2019). This situation has arisen because the residues of the sSgII-70 polypeptide that went into the Edman machine were not read in the correct order, except for an initial M, but were instead mapped in 3D space, antibody like, to produce the EPL001 output sequence. So, an antibody raised to an EPL001-sequence peptide sees the sSgII-70 input sequence. Why would the Edman machine misread the input sequence? Because sSgII-70 depolymerises progressively in the reaction chamber, displaying residue α-amines out of order in a relatively reproducible manner, for the Edman machine to read them faithfully (Hart et al, 2022).

The endogenous epitope of the main antibody raised against the synthetic 14mer peptide EPL001 is analysed to be the SgII-related discontinuous KE·F·NNI, as just described. Residues at both ends of the SgII proteoform are seen simultaneously by the antibody (Fig. 11). For these remote residues to be adjacent SgII-70 must be non-linear. The active face conception and successful mimesis using short synthetic peptides also implies juxtaposed ends (Fig. 11). In ovine SgII-70 the deduced active face is **<u>M·KPVF·N</u>**, with five residues from the N terminus plus N, as in N68, from the C terminus. ‘Adjacent termini’ is a structural prediction for sSgII-70 of the Fig. 10 magic grid. The aa data indicate a non-linear reading path for the Edman sequencing, taking in sSgII-70’s N terminus (with initial M) and its C-terminal NNI. Also suggestive of molecular end-adjacency is the ‘Extension’ part of the EPL001 Extension sequence, which after EPL001’s C-terminal FNNI proceeds for seven further cycles (S1 Table 1). On the assumption that residues D, I & Y derive from the C terminus, there is an apparent shuttle between the two ends of SgII-70, with N-terminal contributions shaded and C-terminal placements underlined: **K**/I **G F**/<u>Y</u>/<u>D</u> x **F**/**V** I/**V** I. ‘Both endedness’ is also indicated by a mixed signal in Harwell 1, **V**/I, while ‘Harwell 2’ of S1 Table 1, in its 8-reading entirety, is an interleaved sequence: **M** <u>N</u>/**F P**/I/M **L** <u>N</u> **V**/<u>A</u> I **T**/**P**.

The Edman reading paths for EPL001 and related sequences can be represented as spirals (S1 Figs. 1 & 2). In the magic grid of Fig. 10 the EPL001 slalom zigzags and the sSgII-14 peaks and troughs likewise evoke spirality. The grid’s first EPL001 switchback sees a reading of SgII-14’s P8 followed by a reading of its L2. A similar switchback is evident in all the ovine Edman sequences, but this effect is not exclusively associated with proline, as in two cases the apex of the switchback is SgII-14’s V9. The ‘pick’ order of EPL001 Edman residues depends on α-amine availability but also steric hindrance, with the latter itself yielding to a spiral conceptualisation (Hart et al, 2022). Spirality is connoted by the maximum-purity minimum-sequence Beale 4 (xx**<u>P</u>**xxxx**<u>V</u>**/_L_xx**<u>F</u>**/_K_**<u>N</u>**xx, S1 Table 1), where _7_**<u>PVF</u>**_9_ is a contiguous triplet motif within sSgII-70. The Beale 4 ‘under signals’, L & K, refer to L2 & K3 within sSgII-70, as in M**<u>LK</u>**TGEKPVFK… The presence of these subsidiary signals is explicable in terms of the spiral conceptualisation just referred to. L2 should be taken second but appears fourth in EPL001 because of steric hindrance by P8. Once P8 is out of the way L2 is available to be read. Removal of V8 releases K3 into availability. The likelihood of the dual positional pairing V/L & F/K arising by chance is 1 in 167 (P = 0.006, significant: Hart et al, 2022). Spirality is also implied in regard to EPL001/sSgII-14 by the playing-card riffle shuffle analysis (see ‘Likely SgII related’). Spirality on a whole-molecule scale is connoted by the seven available ovine N-terminal sequences (S1 Table 1), with all Edman residues accounted for in terms of sSgII-70 at a coverage of 25/70 (36%), as follows, with placements on the assumption that readings D, Y & A are from the C terminus:

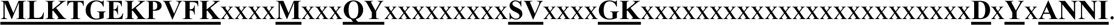

In line with full-molecule spirality is the multi-hundred stepped grand fragmentation pattern detected in MALDI (e.g. Fig. 4, lower panel). Sorting domains within SgII are helical and amphipathic (Courel et al, 2008). Three helical regions are predicted (Fig. 11) for the long centre section (R12-T58) of sSgII-70 not represented on the magic grid of Fig. 10. In sSgII-70’s second sorting domain there is a repeating pattern of hydrophobic residues (shaded) between charged residues (underlined): **ML**KT**G**EK**PVF**K. Amphipathicity of this kind presumably accounts for the bimodalism seen with sSgII-70 in anionex and RP-HPLC, which cannot be due just to a basic N-terminus and an acidic C-terminus, as bimodalism is evinced by Grand Fragments of sSgII-70 in RP-HPLC down to **19mers** lacking the C-terminus altogether (S3 Fig. 3). Coiled coil proteins are prone to homodimerization, another feature of the current data. Summarising, there are multiple reasons for deducing tertiary structural spirality in SgII-70 in terms of both ‘Terminal Spiralisation’ (Fig. 10 grid) and ‘Extra-terminal Spiralisation’ (off-grid), as there are multiple separate reasons for discerning juxtaposed termini.

Treat as a black box the intracellular processing of secretogranin II, including the disassembly of a possible sorting super-domain. Into the black box goes SgII (modularised in Fig. 9 as ABCDE) and out of it comes the SgII-70 proteoform. Four transformations have occurred in the black box to produce SgII-70: (i) *excision* of 61mer and 9mer aa sequence modules B & D from one or two molecules of SgII; (ii) *splicing* of these sequence modules in the order 9+61 (DB) to provide a 70mer polypeptide; (iii) *juxtaposition of termini* and (iv) *spiralisation*.

On (iii), juxtaposition of termini, said termini are in any case anomalous in that the N terminus is Edman baffling, associated with purification bimodalism, displays water losses in high degree and provides as N-terminal fragments *all* SgII-70 ions detected in MALDI, while the C terminus contributes NNI anomalously to an EPL001 Edman sequence which is supposed to be ‘N terminal’. In regard to end-adjacency a molecular hairpin has been posited, speculatively stabilised and twisted by crosslinks (Hart, 2021). By way of comparison, organotrophic IGF-1 has two intramolecular disulphide bonds, insulin three. Yet there are none of the required cysteine residues in the inferred primary structure of (antiorganotrophic) SgII-70 and indeed lack of disulphide bonds is a secretogranin feature. An alternative explanation of SgII-70’s behavioural oddities in MALDI and Edman could be intramolecular crosslinks between aa side chains in the form of isopeptide bonds. These are 8-backbone-atom covalent transamidation crosslinks typically between the side chains of glutamines and lysines. Such Q-K bonds are found in thyroglobulin, the precursor of thyroid hormones, and in the clotting protein fibrin (Walsh, 2006, Fuentes-Lemus et al, 2021). Protease resistant isopeptide stabilization is important in brain function, for example, and skin strength. sSgII-70 is protease and heat resistant, the latter in particular another secretogranin feature. The presence in SgII-70 of isopeptide bonds would not greatly disrupt the MS datafitting of Table 1 as at issue would be reductions for NH_3_ of 17.031 instead of H_2_O at 18.015 Da. Such a datafitting has been attempted for two isopeptide QK bonds within sSgII-70 (Hart, 2021), but exhaustive molecular modelling in silico has failed to deliver an image of sSgII-70 with juxtaposed termini (S5). Additional isopeptide bonds can be contemplated, but the aberrant ions that might be expected from a crosslinked polypeptide fragmenting in MS have not been seen. (There is also no evidence either of additive post-translational modifications, including glycan and lipid attachments.) Crosslinking is not implied by the ‘linear polypeptide’ tryptic digest analysis. Yes, sSgII-70 is protease and heat resistant, but the stability is evidently ‘cliff edge’ and therefore inconsistent with covalent crosslinking, granted apparent partial disintegration in MALDI and Edman. Covalent crosslinking does not explain Edman Nonsequentialism, according to a simulation. Using sSgII-14 (= EPL143: Hart et al, 2022), an E and a K were bonded to the side chains of K and E, respectively. The Edman machine classed the cleaved backbone residue connected to a substituent amino acid as eluting anomalously, registering an x in the correct position in the sequence (S2). *The conclusion is that there are no covalent crosslinks within SgII-70.* An elusive factor was indeed only trapped as a result of an IP/LC-MS antigen capture campaign through *forced* crosslinking using formalin fixation, followed by antigen retrieval.

Transforming sSgII-14 into the EPL001 Edman result requires the movement of only five sSgII-14 residues, four charged and one in a class of its own: K3**^+^**, K7**^+^**, K11**^+^**, E6^−^ & P8 (uniquely a proteinogenic *secondary* amino acid):

The probability of choosing one uncharged residue of any kind from sSgII-14 and the four charged residues is 10/14 x 4/13 x 3/12 x 2/11 x 1/10 x (5!/(4! x 1!)) = 5/1001 or about 1 in 200 (P = 0.005, significant). The probability of choosing proline specifically as the uncharged residue, together with the four charged residues is 1/14 x 4/13 x 3/12 x 2/11 x 1/10 x (5!/(4! x 1!)) = 1/2002 or about 1 in 2000 (P = 0.0005, significant). So charged residues are very much at issue. The Edman EPL001 reading seems to have arisen because of aberrant availability of free amine groups due to out-of-sequence peptide bond cleavages affecting charged residues, amounting to a multistage (non-enzymatic) depolymerisation. Charged residues seem important even by omission, with Ks and Es absent from the First Sighting & Harwell 2 (S1 Table 1). The theme of charged residues continues with the EPL001 Extension, where the only EPL001 residue lacking is E, with a blank x registered instead. Of 14 cycles involved in the production of the EPL001 Edman sequence, four are associated with peptide bond cleavages relating to ‘anomalous lysines’ (Hart et al, 2022).

Of sSgII-70’s residues 25 are charged, representing over a third of the complement. There are ten basic residues, 14% of the total. Four are near the N terminus (K3, K7, K11 & R12), with the rest in the second half of the molecule (K38, K46, H47, R49, K55 & K66). The ten basic residues are outnumbered by 15 acidic Ds & Es (21% of total residues), meaning that sSgII-70 is acidic overall, confirmed by its behaviour in anionex and consistent with high solubility in water (S1 Fig. 11). The acidic residues are distributed thus in the two halves of the molecule, with polyanionic clusters towards the C terminus: _6_ExxxxxxxxExxEExxxxxxxxxxExxxxxE_35_ & _48_ExxDEExxxxxDDEDD_63_. The N terminus is basic, the C terminus acidic, as remarked earlier. ‘Charged residues’ evokes salt bridging. Such bonds are *not* predicted in the molecular modelling of a linear 70mer, the energy minimization process taking into account changes primary, secondary (hydrogen bonding) and tertiary (including ionic bonds, i.e. salt bridges). This does not preclude the possession of such bonds by SgII-70 emerging from the SgII processing black box. Salt bridges are non-covalent electrostatic bonds that arise most commonly from the anionic carboxylate (RCOO^−^) of either D or E and the cationic ammonium (RNH_3_^+^) of K or the guanidinium (RNHC(NH_2_)_2_^+^) of R. Both the charged residues of the salt-bridge can form a hydrogen bond to the same adjacent glutamine residue, providing additional bracing stabilization in the form D/E^Q^K/R (Ruprecht et al, 2014). The involvement of Q in the present case is suggested by the Edman sequencing data of S1 Table 1. A central-sequence motif in EPL001, MKPL**<u>T_5_GKV_8_</u>**KEFNNI, finds echoes in other sequences in the Edman series: **<u>_5_T/GKV_7_</u>**, **<u>_5_TGQA_8_</u>**, & **<u>_4_T/GQY_6_</u>**. In the event of sSgII-70’s K3 (Fig. 10) being read, NNI was read later, while sSgII-70’s Q20 was the start of excursions: **TGQAMEF** (where A67 seems to have been read next) and **T/GQYS/VG?K?** (where the doubletons TG**·**QY**·**SV**·**GK are found in the correct order within sSgII-70, furnishing enough information for a spiralised Edman reading path to be discerned, supporting both Terminal and Extra-terminal Spiralisation: S1 Fig. 2). These perambulations exemplify Edman Nonsequentialism, which, to repeat, is an observation not a conjecture.

Running the C terminus of sSgII-70 up the 45° diagonal of the magic grid from _68_NNI_70_ at bottom right aligns _59_DDEDD_63_ with _3_KxxxK_7_ of the N terminus, suggestively in regard to salt bridging (Fig. 10). E6 and K66 also potentially align. Such interactions could account for Terminal Spiralisation. What of Extra-terminal Spiralisation? That part of the molecule not represented in the magic grid, R12-T58, cannot be an uncomplicated circlet. Physical modelling (i.e. manual construction of a paper tape representation, with the 70 aa marked along it in the single-letter code) suggests a sole twist is likely, with two twists a lesser possibility, even three. This means that the molecule probably resembles a minimally braided (single twist) molecular ampersand, &, with entwined ends. SgII-70 can thus be speculatively envisaged as an amphipathic polypeptide featuring a polyacidic D/E tail looped back to a basic N terminus, bringing about electrostatic self-entanglement, with the rest of the molecule subject to a figure-of-eight spiralisation, the knotty totality – displaying in all two or more ‘crossings’ (Lim & Jackson, 2015; Michielotto, 2024) – potentially accounting for protease and heat resistance and triangulated recalcitrance in MALDI, Edman and molecular modelling in silico.

Endogenously, the prohormone SgII, like granins generally, is self-aggregating in conditions of high calcium and acidic pH, such as are found in the intragranular milieu. The main predicted aggregation sites in the sSgII sequence of Fig. 9 are the N-terminal _18_LIFLL_22_ and the C-terminal _585_LLVKVLEYL_593_ (Tango). These aggregation sites are in modules A & E of Fig. 9. (No aggregation sites are predicted for sSgII-70, consistent with the factor circulating as a monomer, free of artefactual dual decrementation.) Molecules of sSgII can be imagined thus, using the Fig. 9 modularisation: A1B1C1D1E1 + A2B2C2D2E2 + A3B3C3D3E3 + A4B4C4D4E4 etc. In a simple chain arrangement, the aggregation sites would be E1A2, E2A3, E3A4 and so on. Suppose the 9mer (module D) and 61mer (module B) of sSgII-70 come from different molecules of sSgII, a more plausible idea than reverse splicing of DB from a single molecule of SgII. The adjacent splicings to produce sSgII-70 from a chain could therefore be D1B2, D2B3, D3B4 and so on. At issue could be the processing of an aggregated and probably helically entwined sorting super-domain, accounting for the knotted shape of sSgII-70.

Compensatory renal growth in a rat model in vivo involving unilateral nephrectomy was significantly reduced by Candidate 7500 (Hart, 2001) and dose-dependently by EPL001 (Haylor et al, 2009). Crucial to this model is an increase in the bioactivity of endogenous IGF-1, the growth-promoting mitogen, most probably from the systemic circulation (Haylor et al, 2000). An anti-EPL001 antibody administered on its own caused compensatory renal growth to overshoot (Haylor et al, 2009). This is consistent with the neutralisation of an endogenous inhibitor and strengthens the case for an inverse relatedness in this system between EPL001 associated inhibition and IGF-1 stimulation. An extended version of EPL001 is EPL120, having the aa sequence MKPLTGKVKEFNNIKGFGVI (derived from the Edman Extension of S1 Table 1). EPL120 showed dose-dependent inhibition of the (exogenous) IGF-1 stimulated growth in cell numbers of MCF 7 human breast cancer cells over 72h across the EPL120 dose range 0.04 nM - 4 µM (Hart, 2008). Similar results have been obtained with EPL001, albeit with more variable evocation. The superior bioactivity of the EPL120 20mer over the EPL001 14mer, though they share a string of 14 residues, has been explained topologically within a concept of bimodular tri-residue attachment to a cellular receptor which is itself dimeric (Hart et al, 2022). The forward module, MKP, of the sSgII-70 active face, **<u>M·KP</u>**VF·N, displays equal accessibility as represented in the two peptides, for binding to the receptor, but EPL001’s rearward module, VFN, has inferior accessibility to the identically positioned VFN in EPL120. A molecular quirk, this, of adding six residues to EPL001. In the kidney research, EPL001 did not affect IGF-1 stimulated proliferation of mouse collecting duct kidney cells in vitro (Haylor et al, 2009). This indicates that when EPL001 and related peptides inhibit cell proliferation the mechanism does not involve IGF-1 receptor antagonism. It is envisaged that micrin/SgII-70, being antimitogenic and cell-size-limiting, is the (stoichiometric: S2) inhibitory counterpart of IGF-1 within the global organ-size sensing system, via an IGF-1 independent mechanism. The mammalian data just cited are consistent with the findings in nematodes, where the equivalent of IGF-1 is DAF-2. EPL001 and related peptides prolong worm lifespan and modulate fecundity (Davies & Hart, 2008; Davies et al, 2015), as well as altering innate immunity (Davies et al, 2008), all DAF-2 associated aspects of metabolism.

Antiorganotrophic micrin/SgII-70 is thus conceived as the counterpart of organotrophic IGF-1. Both are polypeptides of 70 aa. The former is acidic, the latter basic, with predicted isoelectric points (ExPASy Compute pI/Mw) of 4.64 for hSgII-70 and 7.76 for hIGF-1 (P05019). The bioactivity of SgII-70 is conceptualised to involve, like IGF-1, a dimerized receptor (Hart et al, 2022). IGF-1 has two intramolecular disulphide bonds, as described earlier. Compared with IGF-1, the generality of bioactive peptides (having cell-cell influence) are smaller (6-60 aa), are processed from shorter preprohormones (50-300 aa) and have fewer crosslinks (based on 0, 2 or 4 cysteine residues) (Kastin, 2013).

Even though Candidate 7500 turned up obligingly as part of a precipitate on a 3 kDa filter membrane, the yield was low (Fig. 4 legend), as multiple failed attempts at chemical sequencing attest. ‘Cliff edge’ lability makes SgII-70 resistant to bulk purification – a desirable goal for structural substantiation – and it is currently impossible to provide the factor in true-to-life synthetic or expressed form by dint of apparent topological complexity. Yet it has proved possible to simulate aspects of SgII-70’s proposed role in the organotrophic system using stand-in molecules, crafted on the basis of deduction and ready for exploitation. Therapeutic uses for these hexapeptide effectors include prostate shrinkage in BPH (S2) and reversal of tissue overgrowth in endometriosis, PCOS and cancer, with infertility among potential reproductive applications.

The suggestion that there is an inhibitor of tissue masses in the form of an ‘enigmatic ovarian influence’ (Hart, 1990a) led to a theoretical conceptualisation of the body’s hormonal brake against tissue overgrowth, dubbed micrin (Hart, 2014). This prompted the identification of a candidate prohormone, the neuroendocrine protein secretogranin II (Hart et al, 2017), and a hormonal proteoform of that, SgII-70, enabling the development of hexapeptide mimetics (Hart et al, 2022). Micrin/SgII-70 is conceived to be an antiorganotrophic humoral factor acting remotely in the manner of a classical hormone, with traditional ductless gland (gonadal) provenance and stimulated by antioestrogens (Hart, 2014). SgII-70’s granin stoichiometry is suggested as the counting mechanism behind the regulation of tissue masses. The internal-size-regulating organotrophic system is viewed as having as a key element the hypothalamic-pituitary-gonadal axis, with adrenal input. It is hypothesized that micrin braking, gonadal and hypothalamic, is lifted at puberty to facilitate maturation and wanes with age, bringing on prostatic enlargement and cancers (S6). It is predicted that the endogenous hormone will reverse the pituitary hypertrophy that follows castration in the rat. This effect was attributed in the 1930s to the absence of non-steroidal testicular ‘inhibin’ (McCullagh, 1932), before that name attached itself to modern-day inhibin. The latter polypeptide hormone, although gonadal, acts irrelevantly at the pituitary tier to downregulate follicle-stimulating hormone, rather than at the hypothalamus to suppress the pituitary. Micrin/SgII-70 is McCullagh’s inhibin of yore.

## Supplementary Information

Figshare: Supplementary Information 1: Candidates & Purifications. https://doi.org/10.6084/m9.figshare.27110071.v1. This project outlines the candidate protein and purification techniques utilised.

Figshare: Supplementary Information 2: Assay Methods and Results. https://doi.org/10.6084/m9.figshare.27110170.v1. This project details the methodology used in exemplifying antiorganotrophism for the candidate molecule.

Figshare: Supplementary Information 3: Mass Spectrometry Studies of Candidate 7500. https://doi.org/10.6084/m9.figshare.27110284.v1. This project presents MALDI-TOF MS data of the polypeptide Candidate 7500 and describes it as a novel proteoform of the neuroendocrine prohormone secretogranin II, dubbed sSgII-70.

Figshare: Supplementary Information 4: Tryptic Digests. https://doi.org/10.6084/m9.figshare.27110311.v1. This project attempts to elucidate the identity of Candidate 7500/sSGII-70 using trypsinisation.

Figshare: Supplementary Information 5: Molecular Modelling of sSgII-70. https://doi.org/10.6084/m9.figshare.27110320.v1. This project presents in silico molecular modelling of sSgII-70 directed at a conformational understanding of the molecule.

Figshare: Supplementary Information 6: Extended Discussion. https://doi.org/10.6084/m9.figshare.27110332.v1. This project provides a discussion of possible chains of logic and associated theoretical frameworks which can be associated with and possibly explain the generated organotrophic data and MALDI analyses.

Figshare: Supplementary Information 7: Arrive 2.0 Guidelines. https://doi.org/10.6084/m9.figshare.27110371.v1. This project provides the Arrive 2.0 Guidelines.

## AUTHOR CONTRIBUTIONS

The authors confirm that they are the originators of this work and that they have approved it for publication. The views expressed are theirs alone. JEH was lead author. Besides contributing editorially, authors had these special responsibilities: JEH (project management, data analysis, statistics), KGD (nematology, reproductive studies), CRM (bioinformatics), ACH (MS data reanalysis), DRH (IHC, graphics) & RPN (MS, molecular modelling, internal peer review).

## ACKNOWLEDGEMENTS

For personal communications the following are lauded and thanked: Pat Barker, Carolyn Carr, Clare Collier, Ed Dudley, Ben Ferneyhough & Julian Hiscox. Maureen Hamon’s input is appreciated on the work at Babraham, where ligand purification was pioneered by Dennis Beale. Factor hunters thereafter included Andy Scutt, Charles Dickerson & Dave Emery. Dirk Winkelhardt is thanked for tryptic digest database assistance. Gratitude is expressed to Mark Silverman for help with the riffle shuffle analysis. JEH salutes all the researchers who have been involved in the project and the investors in Endocrine who made the work possible.

## COMPETING INTERESTS

JEH is founding scientist of Endocrine Pharmaceuticals of Hampshire, UK (Company No. 03005721) and is an employee of the company, holding shares and share options. Endocrine has relevant patents and patent applications which in no way constrain sharing of data and materials. All coauthors are participants in Endocrine’s share option scheme and most of those mentioned in Acknowledgements hold shares or share options in Endocrine.

## FUNDING

The early research benefitted from a SMART Feasibility award (YHF/21865/SM00) and a SMART Development award (YHF/21865/SM02) from the UK government. No other specific grant has been received from any other funding agency in the public, commercial or not-for-profit sectors. The research was otherwise paid for by Endocrine Pharmaceuticals of Hampshire, UK (Company No. 03005721).

## ETHICS

All experimental procedures, including those in vivo subject to the research checklists of S7, were performed in compliance with applicable laws, regulations and professional standards of probity and good faith, with appropriate ethical oversight. The studies described in S2 were conducted under United Kingdom Home Office Project Licence Nos. PPL 80/1025, PPL 80/1150 & PPL 30/1487. As few animals were used as possible, in accordance with ‘Reduction’ in the Three Rs of Replacement, Reduction, Refinement. In the matter of sample identity, proprietary codes were favoured over explanatory designations to foster blind experimentation with unbiased analysis and impartial supervision. This objective was aided in any case by project inscrutability, which countered confirmation and scepticism bias. Protocols were pre-registered to comply with UK Home Office licencing requirements and for ethics submissions but were not pre-registered in an open database, being exploratory studies intended to culminate a hormone discovery project involving prior publications, wherein are described diverse materials (with provenances) and methods (with protocols and permissions). Data integrity has been upheld throughout, with no exclusion of outliers, no withholding of null findings and with appropriate recording and archiving. Raw data are available via supplementary files S1-6, in the interests of transparency and the provision of a true account.

## Notes

https://figshare.com

